# Cell type-specific epigenetic regulatory circuitry of coronary artery disease loci

**DOI:** 10.1101/2025.02.20.639228

**Authors:** Dennis Hecker, Xiaoning Song, Nina Baumgarten, Anastasiia Diagel, Nikoletta Katsaouni, Ling Li, Shuangyue Li, Ranjan Kumar Maji, Fatemeh Behjati Ardakani, Lijiang Ma, Daniel Tews, Martin Wabitsch, Johan L.M. Björkegren, Heribert Schunkert, Zhifen Chen, Marcel H. Schulz

**Affiliations:** Department of Medicine, Institute for Computational Genomic Medicine, Goethe University Frankfurt, 60590 Frankfurt, Germany; Deutsches Zentrum für Herz- und Kreislaufforschung (DZHK), Rhein-Main, Germany; Department of Cardiology, German Heart Centre Munich, School of Medicine and Health, Technical University of Munich, 80636 Munich, Germany; DZHK, Partner Site Munich Heart Alliance, Munich, Germany; Department of Genetics & Genomic Sciences, Institute of Genomics and Multiscale Biology, Icahn School of Medicine at Mount Sinai, New York 10029, USA; German Center for Child and Adolescent Health (DZKJ), Partner Site Ulm; Division of Pediatric Endocrinology and Diabetes, Department of Pediatrics and Adolescent Medicine, Ulm University Medical Center, 89075 Ulm, Germany; Department of Medicine, Karolinska Institutet, Karolinska Universitetssjukhuset, 14157 Huddinge, Sweden

**Keywords:** Coronary artery disease, epigenetics, genome-wide association studies (GWAS), Gene-Based Association Test (GATES), SNP exploration and analysis using epigenomics data (SNEEP), CRISPR/Cas9, and non-coding RNA genes

## Abstract

Coronary artery disease (CAD) is the leading cause of death worldwide. Recently, hundreds of genomic loci have been shown to increase CAD risk, however, the molecular mechanisms underlying signals from CAD risk loci remain largely unclear. We sought to pinpoint the candidate causal coding and non-coding genes of CAD risk loci in a cell type-specific fashion. We integrated the latest statistics of CAD genetics from over one million individuals with epigenetic data from 45 relevant cell types to identify genes whose regulation is affected by CAD-associated single nucleotide variants (SNVs) via epigenetic mechanisms. Applying two statistical approaches, we identified 1,580 genes likely involved in CAD, about half of which have not been associated with the disease so far. Enrichment analysis and phenome-wide association studies linked the novel candidate genes to disease-specific pathways and CAD risk factors, corroborating their disease relevance. We showed that CAD-SNVs are enriched to regulate gene expression by affecting the binding of transcription factors (TFs) with cellular specificity. Of all the candidate genes, 23.5% represented non-coding RNAs (ncRNA), which likewise showed strong cell type specificity. We conducted a proof-of-concept biological validation for the novel CAD ncRNA gene *IQCH-AS1*. CRISPR/Cas9-based gene knockout of *IQCH-AS1*, in a human preadipocyte strain, resulted in reduced preadipocyte proliferation, less adipocyte lipid accumulation, and atherogenic cytokine profile. The cellular data are in line with the reduction of *IQCH-AS1* in adipose tissues of CAD patients and the negative impact of risk alleles on its expression, suggesting *IQCH-AS1* to be protective for CAD. Our study not only pinpoints CAD candidate genes in a cell type-specific fashion but also spotlights the roles of the understudied ncRNA genes in CAD genetics.

## 1 Introduction

The past 15 years have witnessed an explosion of genetic discoveries on common complex diseases, mainly by genome-wide association studies (GWAS). For coronary artery disease (CAD), > 300 gene loci have been associated with genome-wide significance^1–3^, deepening the understanding of disease pathology. Downstream to the genetic discoveries, a bigger challenge is to fill the gap between genetic variants and disease risk. To tackle this, functional studies are indispensable to ultimately unveil related molecular mechanisms and inform novel diagnostic and therapeutic developments. Importantly, GWAS signals are most often found in non-coding regions of the genome, which indicates a major role for altered gene regulation in the etiology^4^. In this respect, downstream causal genes, and cell types of action need to be elucidated allowing us to pinpoint the biological mechanisms underlying CAD. To date, efforts have been made to prioritize causal genes in CAD loci by various independent or joint methods by assessing 1) biological plausibility, 2) rare coding variant(s) associated with CAD, 3) likely pathogenic variant(s) relevant to CAD in ClinVar^5^, 4) evidence from cardiovascular (CV) drug(s), 5) causality by Mendelian Randomization studies, 6) a protein-altering variant in high LD with the sentinel CAD variant, 7) expression quantitative trait loci (eQTLs) in a CAD-relevant tissue, and 8) CV-relevant phenotypes in knockout mouse models^1,2,6^. However, these methods were not able to prioritize target genes for all the GWAS loci^6^. Among these methods, eQTL analysis contributed the largest number of candidate genes^2,3,7^, yet eQTLs might explain only a small fraction of GWAS heritability plausibly^8^. More diverse functional genomic readouts beyond transcriptomics are urgently needed to identify disease mechanisms. The epigenome regulates gene expression in cells and engages the response of genetic variation to environmental changes. While sporadic studies have explored tissue-level chromatin states or epigenetic marks at CAD loci^1,9,10^, epigenetics of CAD genetics is rather under-investigated.

In the current study, we systematically investigate the cell type-specific epigenetic circuit of CAD genetics by integrating the summary genetic statistics from over one million individuals with the regulatory elements of 45 types of disease-relevant cell types and Hi-C information. We identified candidate causal genes at CAD loci, which were not uncovered by reported methods. Beyond commonly explored protein-coding candidate genes, our analyses allowed extensive examination of the CAD-associated non-coding RNA (ncRNA) genes which were otherwise depreciated due to insufficient sequencing depth in transcriptome-based causal gene prioritization. Furthermore, biological validation was conducted on one of the novel long non-coding RNAs (lncRNA) to confirm the reliability of our analysis and improve our knowledge of CAD mechanisms.

## Results

### Cell type-specific annotation of CAD-associated SNVs

To explore epigenetic and cell type-specific effects of CAD genetics, we obtained the latest summary genetic statistics of CAD GWAS from over one million human subjects^1^ and bulk or single-cell epigenetic data from 45 disease-relevant cell types (Fig. 1A)(Supp. Table 1&2). 47,635 CAD-associated single nucleotide variants (SNVs) (≤ 1% FDR)^1^ and 24,799 SNVs in LD (R^2^ ≥ 0.8) resulted in a total of 72,432 CAD-SNVs included in our analysis (Supp. Table 3). Based on Ensembl Variant Effect Predictor (VEP)^11^, the majority (67.1%) of CAD-SNVs were intronic variants and less than 10% were variants likely with a stronger effect size, such as missense, UTR or splicing region SNVs (Fig. 1B). Beyond the general annotation using VEP, we further investigated the intersection of CAD-SNVs with epigenetics of the 45 cell types, which covered 12 cell lineages including lymphoid, hematological, myeloid, erythroid, endothelial, myogenic, neuronal, adipogenic, fibroblast, hepatic, epithelial, and mesenchymal cells. On average 121,376 cis-regulatory elements (CREs) per cell type were identified and, in total, 1,243,265 CREs from the 45 cell types. On the one hand, we found that 0.71% (N = 8,864) of all CREs contained a CAD-SNV and 0.34% (N = 4,233) a CAD-SNV affecting TF binding (TF-SNV), suggesting that 47.89% of CREs with CAD-SNVs could regulate TF binding (Supp. Table 4). In addition, CREs with CAD-SNVs shared across cell types were more likely to have a TF-SNV than those unique to a cell type. On the other hand, 18.73% of CAD SNVs were located in CREs, 14.49% intragenic, and 4.24% intergenic (Fig. 1C). Among the CAD-SNVs in CREs (N = 13,563), 39.80% were predicted to affect transcription factor binding (N = 5,397) (Fig. 1D). Vascular and immune cells were the leading cell types having relatively large fractions of CRE- and TF-SNVs, such as endothelial cells (ECs), smooth muscle cells (SMC), fibroblasts, T cells, monocytes, and neutrophils (Fig. 1E), in line with their crucial roles in cardiovascular health. 957 TF-SNVs were located in CREs of neurons, resonating with the roles of neuroimmune interfaces in atherosclerosis ^12^.

**Figure 1:**
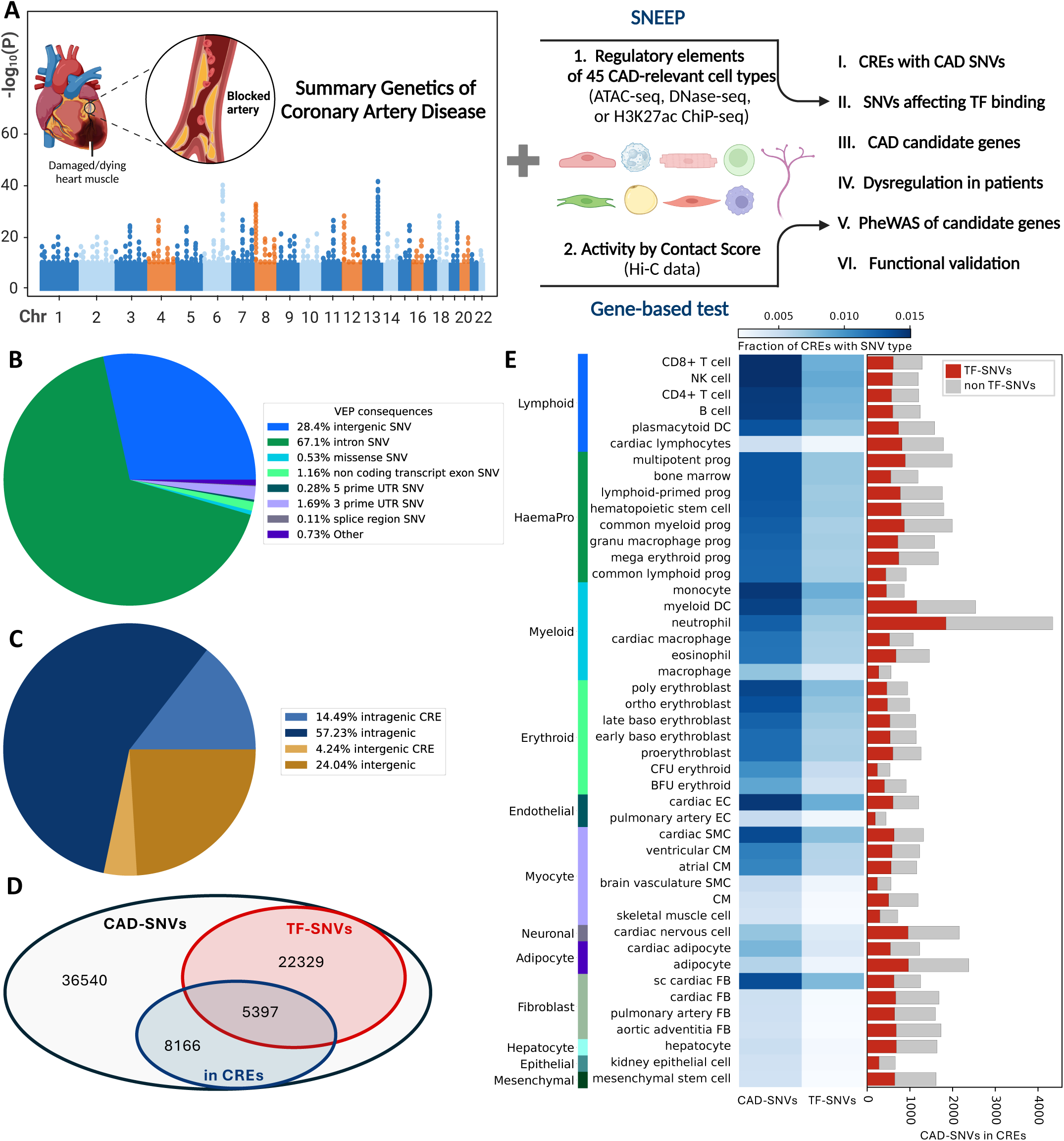
Analysis outline and cell type-specific annotation of single nucleotide variants (SNVs) associated with coronary artery diseases (CAD). **(A)** Overview of the project. GWAS of CAD^1^ was combined with epigenome data and predicted cis-regulatory element (CRE) to gene interactions for 45 cell types. Candidate CAD genes were identified with a tool to annotate transcription factor (TF) SNVs (SNEEP^14^) and a gene-based association test (GATES^13^). The results were supplemented with data of CAD patients, PheWAS analyses and the functional validation of a candidate CAD gene. Created with BioRender.com. **(B)** Functional consequences of CAD-SNVs (p-value ≤ 2.52E-5 and SNVs in linkage disequilibrium) annotated with the Ensembl Variant Effect Predictor (VEP)^11^. Only one consequence is considered per variant. **(C)** Location of CAD-SNVs with respect to genes and CREs. **(D)** Venn diagram of the intersection of CAD-SNVs, TF-SNVs and SNVs located in CREs of any cell type. **(E)** Overlap of CAD-SNVs and TF-SNVs with CREs across cell types and grouped by lineage. Lineages sorted by average fraction of CREs with CAD-SNVs. NK cell: natural killer cell; DC: dendritic cell; BFU erythroid: burst-forming unit erythroid; CFU erythroid: colony-forming unit erythroid; CM: cardiomyocyte; EC: endothelial cell; SMC: skeletal muscle cell; FB: fibroblast.

### Integration of epigenetic data reveals novel CAD candidate genes

Based on the epigenetic data of the 45 cell types (Fig. 1E), we used two complementary approaches to prioritize candidate genes for CAD, namely, the Gene-Based Association Test (GATES)^13^ and the gene prioritization using SNEEP^14^ (Methods). After assigning CREs to their target genes for each cell type^15^, both were applied. We obtained 1,387 genes with GATES and 503 genes with SNEEP resulting in a total of 1,580 candidate CAD genes (Supp. Table 5). We compared our candidate genes with reported CAD genes by other genetic studies and genes identified via GWAS-eQTL colocalization analysis^16^ using genotyped transcriptome data of CAD-relevant tissue types from the Genotype-Tissue Expression (GTEx)^17^ and Stockholm-Tartu Atherosclerosis Reverse Network Engineering Task (STARNET) projects^18^. Out of 1,580 genes, 539 genes overlapped with reported CAD genes and 544 genes with CAD eQTL genes in a total of 782 genes, namely, 49.49% of replication rate on genes from other studies or another method (Fig. 2A, Supp. Fig. 1A&1B). 798 genes were novel CAD candidate genes prioritized by our epigenetics-GWAS integration (Supp. Fig. 1B). Gene set enrichment analysis showed *lipid metabolism*, and *TGF beta, VEGFA,* and *interleukin 11 signaling* to be the top pathways for CAD risk mediated by the 1,580 candidate genes (Fig. 2B, Supp. Table 6), in line with known pathways for the disease. Our analysis also pointed to an ncRNA-related pathway, ‘miRNA targets in ECM and membrane receptors’, for CAD (Fig. 2B), which might represent a novel mechanism for the disease. Our 1,580 candidate genes contained 1,208 protein-coding and 372 ncRNA genes (Fig. 2C, Supp. Fig. 1B). Our epigenetic integration identified the highest fraction of ncRNA genes compared to the other three sources, including the well-known cardiovascular disease (CVD) ncRNAs *CDKN2B-AS1*^19–21^ and microRNA132 (MIR132)^22^ (Fig. 2C, Supp. Fig. 1B&1E).

**Figure 2:**
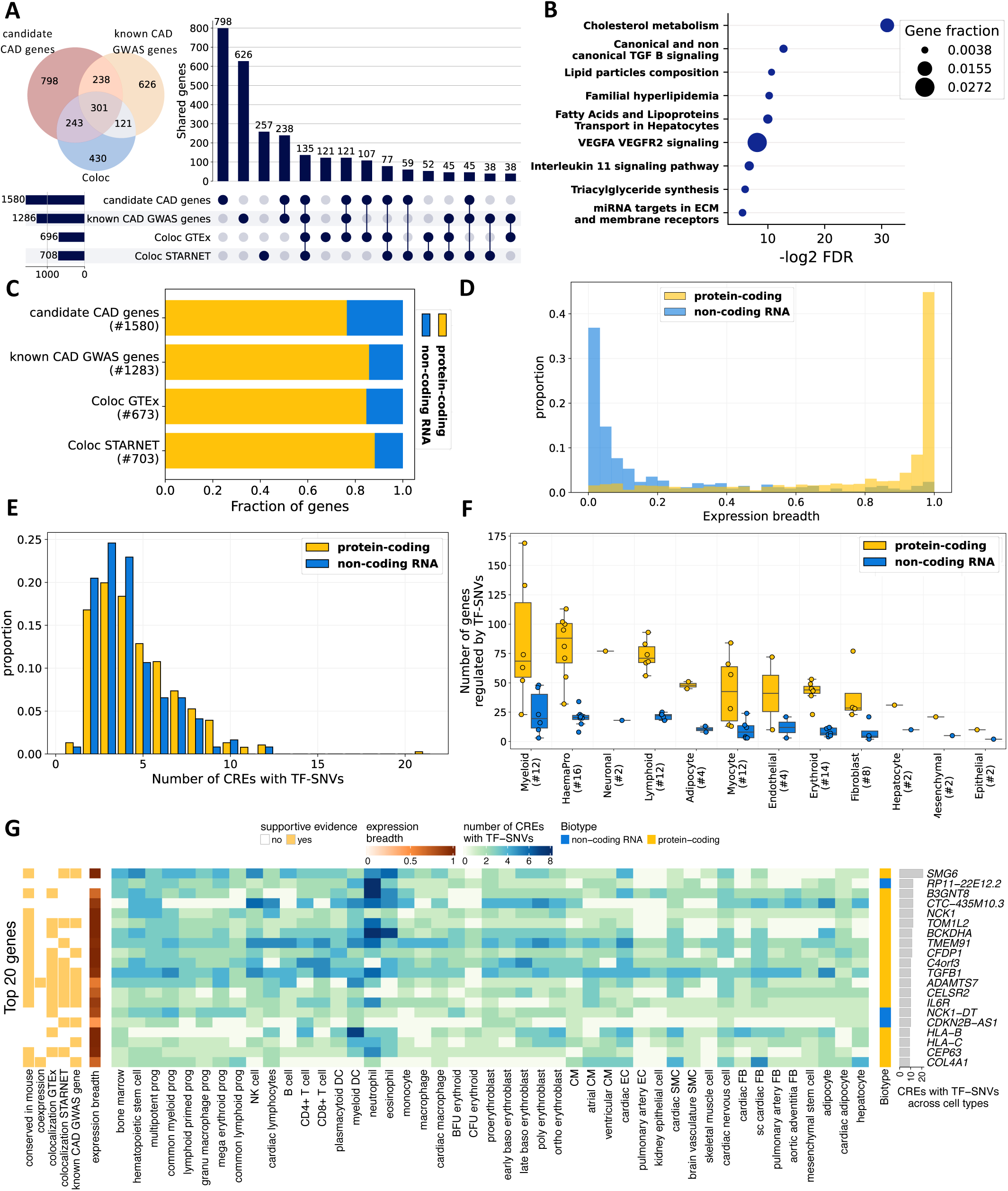
Characterization of candidate CAD genes. **(A)** Intersection of the candidate CAD genes, known CAD loci genes and genes found via eQTL colocalization analysis^16^. In the Venn diagram Coloc refers to the joint set of genes found via GTEx^17^ and STARNET^18^. **(B)** GO term enrichment of the candidate CAD genes. Shown are selected terms from the top 20 enriched terms of the KEGG database (without redundant terms and for Familial hyperlipidemia the adjusted p-value across all five types was averaged). **(C)** Fraction of protein-coding and non-coding RNA genes among the gene sets from **(A)**. **(D)** Expression breadth of the candidate genes, separated by gene biotype. **(E)** Number of CREs per gene that has a TF-SNV, separated by gene biotype. Only genes found via TF-SNVs are shown (N = 503). The CREs with TF-SNV were merged across all cell types in which a gene had a CRE at its promoter and thus was considered active. **(F)** Number of candidate CAD genes identified via TF-SNVs grouped by lineage and separated by gene biotype (boxplot center line mean, boxlimits inter-quartile range, whiskers up to 1.5x inter-quartile range). **(G)** Heatmap of the top 20 genes ranked by the number of merged CREs with TF-SNVs (as described for **(E)**). Additional evidence is added per gene (Supp. Table 5). Per cell type the number of CREs with TF-SNVs is set to zero if the gene is not considered active in the cell type.

To illustrate the expression of candidate CAD genes across different cell types and tissues, we determined their expression breadth in the IHEC EpiATLAS^23^, which measures in how many cell types a gene is expressed (Fig. 2D). As expected, while most of the protein-coding genes were expressed across many cell types, ncRNA genes showed mostly cell type-specific expression. The 503 genes prioritized by SNEEP had an average of 4.5 CREs with TF-SNVs across all cell types (Fig. 2E). The number of identified CAD genes via TF-SNVs varied per cell type (Fig. 2F). Myeloid and haematopoetic progenitor cells had the highest number of genes regulated by TF-SNVs, although the variation was quite large. Surprisingly, many genes (N = 503) were identified via TF-SNVs in different cell types and without a strong indication of genetic linkage, suggesting a range of different regulatory CAD-SNVs linked to the same gene (Fig. 2E).

Among the top 20 genes with the highest number of CREs containing TF-SNV, 10 were known CAD genes, 15 were also identified by the GWAS-eQTL colocalization analysis, and 14 were conserved in mice (Fig. 2G, Supp. Fig. 1D&1E,). The top gene, *SMG6* (Telomerase-binding protein EST1A), had 21 CREs with TF-SNVs mostly in blood and immune cells. *SMG6* has been implicated with CAD through SNVs that affect post-transcriptional RNA methylation^24^, by colocalization of eQTL data^21^ and TF binding analysis^25^. To our knowledge, *B3GNT8* (Beta-1,3-N-Acetylglucosaminyltransferase 8) has not been implicated in CAD yet. However, B3GNT proteins have been associated with diabetes and processes of the immune system^26^, which are risk factors for CAD. One of the three ncRNA genes with a high number of TF-SNV containing CREs is *CDKN2B-AS1*. *CDKN2B-AS1* is located in the first ever identified and still strongest CAD GWAS locus, the 9p21 locus^27–29^. The candidate gene(s) at this locus and their mechanisms of action for CAD have been challenging to investigate^19,20^. We found *CDKN2B-AS1* as a candidate causal gene at this locus with 11 TF-SNVs in 10 different CREs which are mainly active in immune cells (Supp. Fig 2, Supp. Table 15).

### Genetic signals pinpoint CAD-related transcription factors

As a large subset of CAD-SNVs appears to affect TF binding (Fig. 1D), we further studied the TFs involved. The SNEEP-TF test was used to discover TFs that are more often affected by a CAD-SNV than observed by chance (odds ratio ≥ 5, Methods). This analysis was done for each of the 45 cell types of epigenome samples and revealed 38 TFs (CAD-TFs), whose binding was changed by risk alleles of CAD in at least one cell type (Fig. 3A, Supp. Table 7). Among the predicted CAD-TF target genes are protein-coding and ncRNA genes and several are either known CAD GWAS genes or candidate CAD genes. The binding of 26 TFs was affected in more than one cell type. Notably, the binding of the AP-1 family TFs was only enriched for adipocytes, including BATF3, FOSB, and FOS, and the dimers FOS::JUND, FOSL2::JUN, and FOS::JUN. While AP-1 family TFs fulfill a variety of functions, their members have been specifically shown to be involved in differentiation and apoptosis in murine adipocytes^30^. Zhao et al. found that the TFs of the AP-1 family regulate the known CAD genes *SMAD3* and *CDKN2B-AS1*, which are also in our CAD candidate gene set^31^. These data underline the role of adipocytes or adipose tissues in CAD.

**Figure 3:**
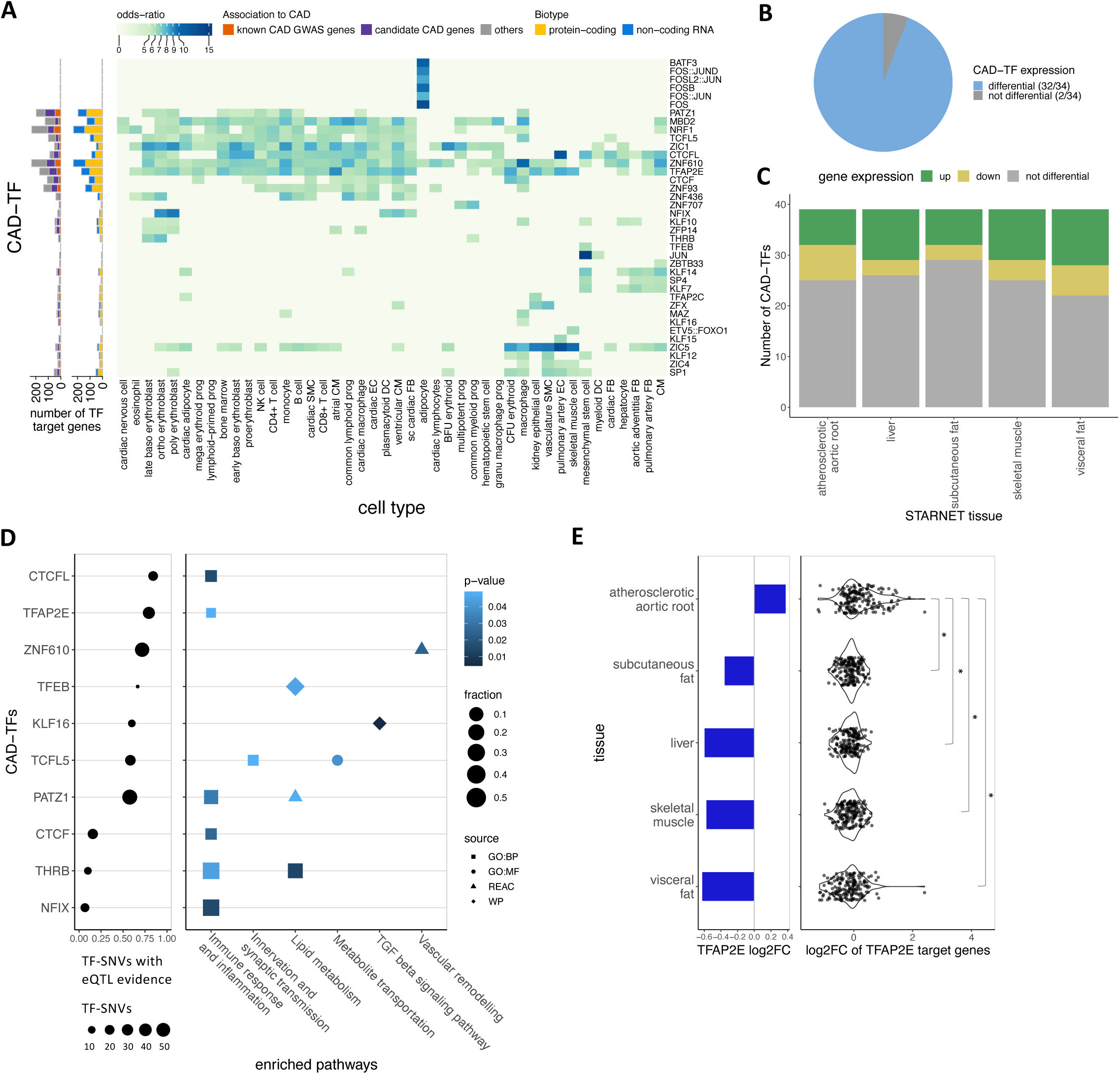
CAD risk alleles affect TF binding. **(A)** Cell type-specific enrichment of CAD-TFs that are affected by binding to TF-SNVs. Each TF regulates a different number of target genes. By the stacked barplots (left) the fraction of target genes associated to CAD and the biotypes are visualized. Analysis of differential expression of CAD-TFs in all **(B)** or individual STARNET tissues^18^ **(C)**. **(D)** CAD-TFs with eQTl support (GTEx^17^) and their associated biological processes and pathways. **(E)** Analysis of TF gene deregulation (log2 fold change, barplot) and expression deregulation of associated TF target genes in different STARNET tissues (log2 fold change, violin scatter) for TFAP2E.

Further analysis using transcriptome data of disease-relevant tissues from controls and CAD patients of the STARNET project^18^ revealed that the majority of CAD-TFs (N = 32) were differentially expressed in investigated tissues (Fig. 3B, Supp. Table 8). Most TFs showed upregulation in CAD-relevant tissues (Fig. 3C). CAD-SNVs regulating TF binding were also identified as eQTLs in CAD-relevant tissues, constituting independent evidence of disease-relevant TFs. The TFs CTCFL, TFAP2E, and ZNF610 had the highest eQTL support and the eQTL genes were involved in pathways for CAD such as *immune response and inflammation*, *lipid metabolism*, *TGF beta signaling*, and *vascular remodeling pathway* (Fig. 3D, Supp. Table 9&10). Interestingly, the eQTL genes of CAD-SNVs regulating TCFL5 binding were enriched for *innervation and synaptic transmission*, a newly discovered pathway for CAD^12^. Further, we observed that for several TFs, the target genes were dysregulated in CAD patients with tissue specificity. For instance, target genes of TFAP2E are upregulated in the atherosclerotic aortic root, but downregulated in fat, liver, and skeletal muscle (Fig. 3E). Similar tissue-specific effects of other TFs were observed as well, including ZIC4, ZBTB33, MBD2, ZNF610, TCFL5 and ZFP14 (Supp. Fig. 3).

### Novel CAD candidate genes associated with cardiovascular risk factors

To investigate the relevance of the identified CAD candidate genes, we performed a phenome-wide association study (PheWAS). The PheWAS datasets were obtained from the Common Metabolic Diseases Knowledge Portal and included the latest GWAS summary genetic statistics of 29 traits of CAD risk factors. We focused on the eQTLs of the candidate genes and extracted the eQTLs from CAD-relevant tissue transcriptomes from GTEx^17^ and STARNET^18^. Based on the probability of PheWAS-eQTL colocalization (Methods), we identified the phenotypic traits bridging our candidate genes and CAD. In total, 1,219 candidate genes, including 1,052 protein-coding and 167 non-coding genes, showed significant colocalization^16^ (PPH ≥ 0.60) with at least one GWAS trait in a relevant tissue type (Supp. Table 11).

Overall, CAD candidate genes demonstrated the highest frequency of associations with inflammatory biomarkers (mainly immune cell counts) followed by lipid levels (mainly cholesterol and triglyceride (TG) levels), both of which involved over 800 genes (Fig. 4A). Artery and adipose tissues, which play essential roles in vascular and metabolic functions, exhibited the highest number of genes with significant colocalizations, with 903 and 806 prioritized genes, respectively (Fig. 4A). By comparison, liver, skeletal muscle, and blood revealed nearly equivalent gene counts with correspondingly 686, 681, and 671 genes. Interestingly, 42 genes showed significant PheWAS-eQTL colocalization signals in kidney cortex tissue, in line with the fact that chronic kidney disease is a risk factor for CAD^32^. For the novel CAD candidate genes (Fig. 4B), we observed similar patterns in comparison to all prioritized CAD genes and known genes (Fig. 4A, Supp. Fig. 4), again, suggesting the reliability of the novel genes.

**Figure 4:**
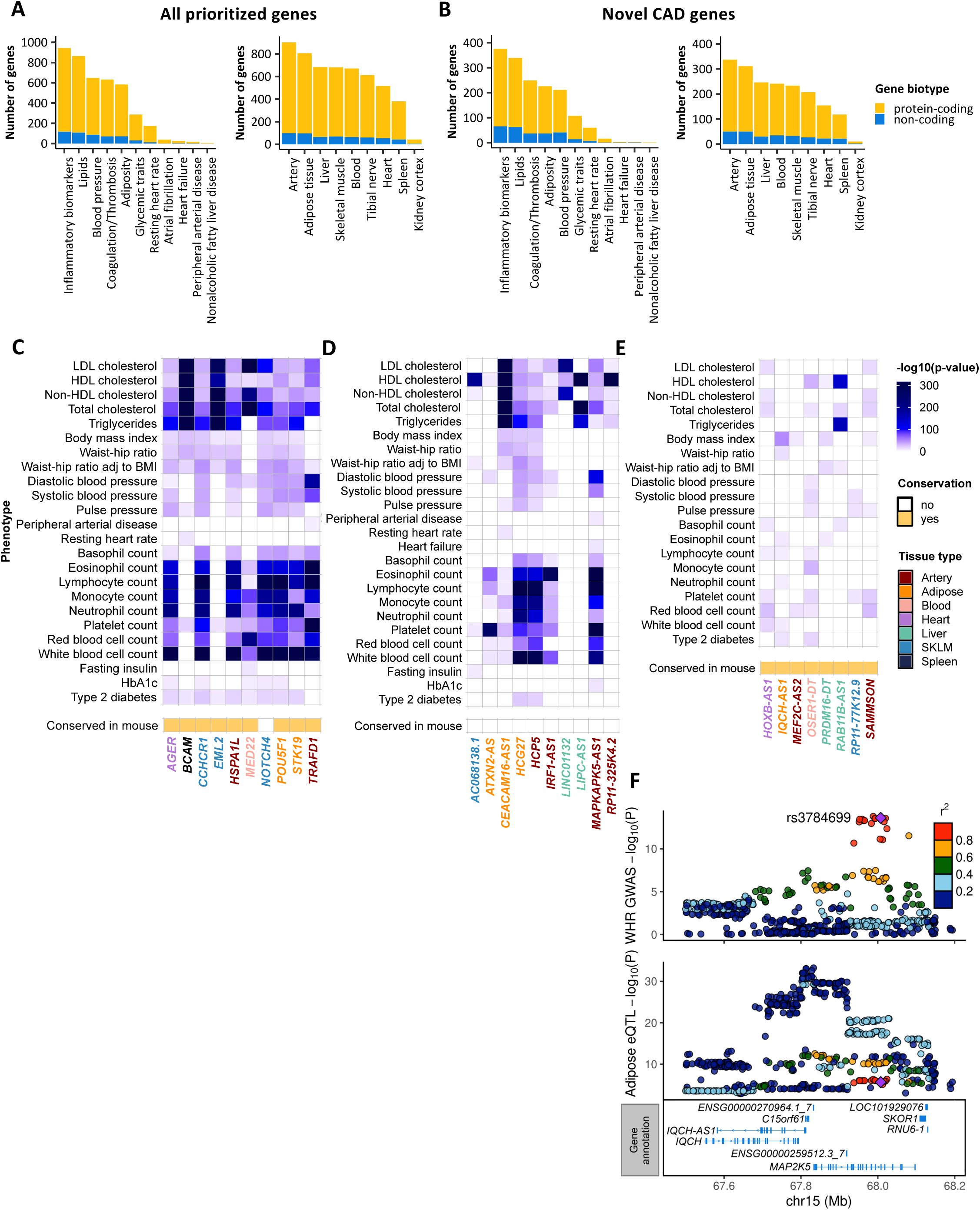
PheWAS-eQTL colocalization analysis on prioritized genes. **(A-B)** Number of protein-coding and non-coding genes with significant colocalized GWAS-eQTL signals across phenotypes and tissue types among all prioritized **(A)** and CAD-novel genes **(B)**. **(C-D)** Top 10 CAD-novel protein-coding **(C)** and non-coding **(D)** genes with the strongest eQTL effects on CAD-related phenotypes. **(E)** Phenotypic association of CAD-novel non-coding genes showed conservation in mouse. **(F)** *IQCH-AS1* locus displaying colocalization of the waist to hip ratio (WHR) GWAS results with eQTL signals of adipose tissue.

For the top novel candidate genes ranked by their association strength with PheWAS traits, both protein-coding and ncRNA genes were predominantly associated with inflammatory biomarkers and lipid levels (Fig. 4C&4D). Our data indicated that ncRNAs could contribute to CAD independent of the host genes, especially for the intergenic ncRNAs, such as LINC01132 and AC068138.1 (ENSG00000235070). These two were also uniquely associated with lipid levels. However, none of the top ncRNA genes showed conservation in mice. Interestingly, among the 167 ncRNA genes with PheWAS-eQTL colocalization signals, eight were conserved in mice (Fig. 4E).

### The novel gene *IQCH-AS1* contributes to CAD via obesity-related phenotypes

To further validate our novel findings, we conducted a biological validation on a novel CAD candidate gene, encoding a conserved lncRNA, *IQCH-AS1*. PheWAS-eQTL colocalization analysis suggested *IQCH-AS1* to be most significantly associated with obesity-related phenotypes including body mass index (BMI) and waist-hip ratio (WHR) (Fig. 4E). The GWAS signals of BMI and WHR were colocalized with *IQCH-AS1* eQTLs from adipose tissue, demonstrating that the roles of *IQCH-AS1* in this tissue might contribute to obesity and therefore increase the risk of CAD (Fig. 4F, Supp. Table 14). We, therefore, studied the *IQCH-AS1* functions in human Simpson-Golabi-Behmel Syndrome (SGBS) preadipocytes and adipocytes. We first generated *IQCH-AS1* knockout (KO) SGBS preadipocyte lines by a dual CRISPR targeting strategy, which used two gene-specific single guided (sg) RNAs to target the shared exon of the major *IQCH-AS1* isoforms *IQCH-AS1*. The dual CRISPR excised 48bp of the exon 13 and dramatically reduced the RNA level of *IQCH-AS1* compared to the scrambled control line (Fig. 5A&5B). By BrdU-based proliferation assay, we observed reduced proliferation of *IQCH-AS1*-KO preadipocytes which might indirectly lead to hypertrophy of existing adipocytes in hyperlipidemic conditions due to reduced source of new adipocytes (Fig. 5C). *IQCH-AS1*-KO preadipocytes had diminished differentiation efficiency indicated by less PPARG expression in the nuclei (Fig. 5D, Supp. Fig. 5). Therefore the corresponding differentiated adipocytes showed less accumulation of TGs (Fig. 5E), which could lead to TG increase in the circulation due to diminished capacity of lipid storage by adipocytes. Furthermore, *IQCH-AS1*-KO adipocytes released more proinflammatory cytokines including pentraxin 3^33^ and IL-18^34^, but fewer anti-inflammatory cytokines including IGFBP-3^35^, Cripto-1^36,37^, VEGF^38^ and IL-10^39^ (Fig. 5F). Our data from *IQCH-AS1*-KO preadipocytes and adipocytes indicates the gene to be protective for CAD. Indeed, *IQCH-AS1* was downregulated in SAT and VAT of atherosclerosis patients from STARNET^18^ cohorts compared to controls (Fig. 5G). Furthermore, the BMI- and WHR-increasing alleles (e.g., rs3784699_C) at this locus were associated with reduced *IQCH-AS1* expression (Fig. 5H&5I). The biological experiments suggest the CVD relevance of our novel gene, *IQCH-AS1*, which further supports the reliability of our epigenetics-GWAS analysis.

**Figure 5:**
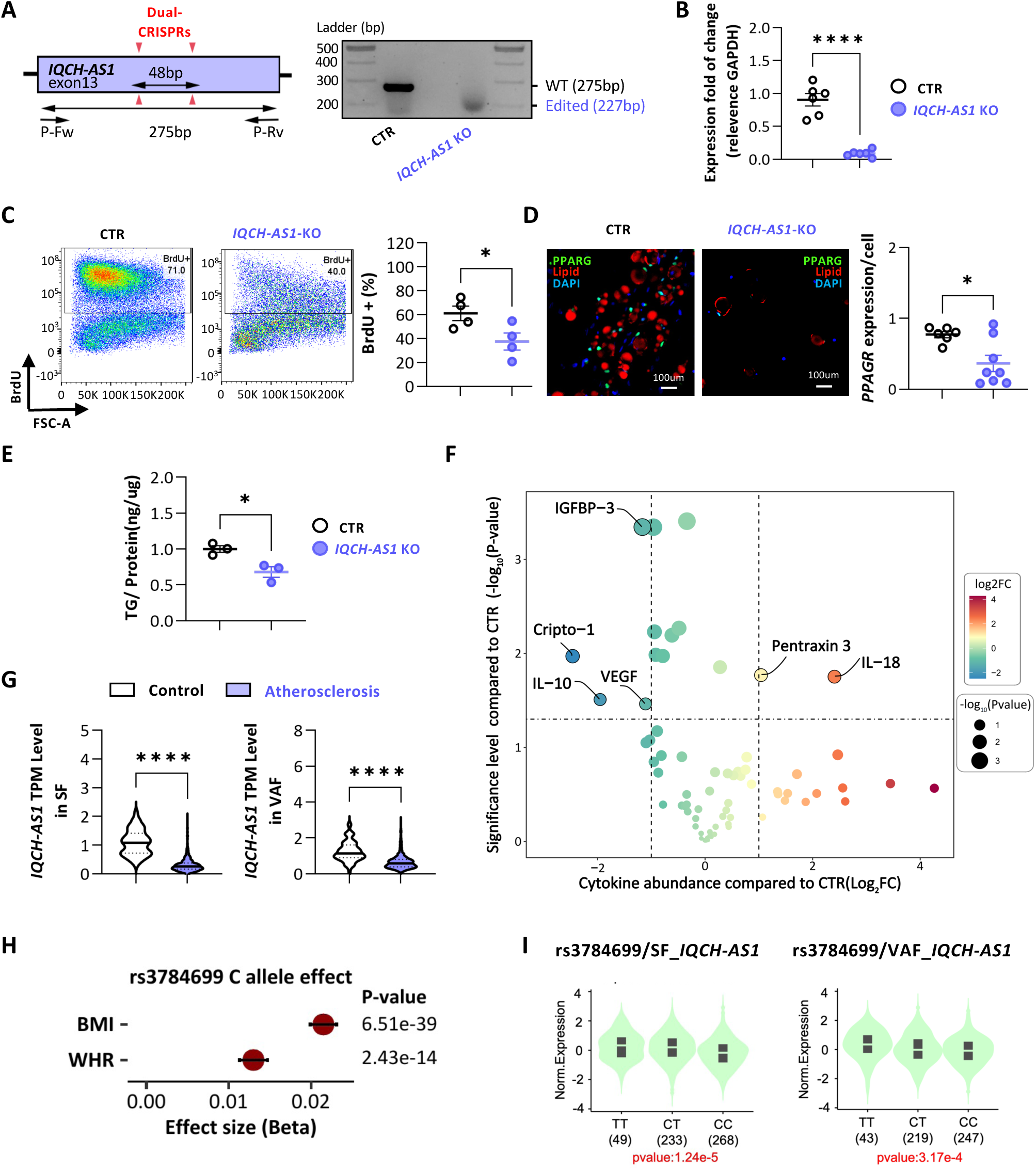
Atherogenic phenotypes induced by CRISPR-based knockout(KO) of IQCH-AS1. (**A-B**) Dual CRISPR-KO of IQCH-AS1 perturbed the gene (**A**) and decreased the RNA levels of IQCH-AS1 (**B**) in SGBS-preadipocytes. (C) KO of IQCH-AS1 reduced the proliferation of SGBS-preadipocytes indicated by fewer BrdU+ cells. (D-E) IQCH-AS1-KO adipocytes had less PPARG expression (**D**) and decreased triglycerides (TG) accumulation (**E**). PPARG was stained by the protein-specific antibody (green). Lipid droplets were labeled by HCS LipidTOXTM Red neutral lipid stain (red), and DNA by DAPI (blue) (Supp Fig. 5). (**F**) Proteome Profiler assay using cell culture medium identified differential cytokine release from IQCH-AS1-KO adipocytes. (**G**) Expression of IQCH-AS1 was reduced in subcutaneous (SF) and visceral abdominal fat (VAF) of atherosclerosis patients (n=568 in SF, n=531 in VAF) in comparison to control individuals (n=92 in SF, n=103 in VAF). Center line is the median, dashed lines upper and lower quartiles. (**H-I**) rs3784699_C (the read SNV in Fig. 4E), was associated with increasing body mass index (BMI) and waist-to-hip ratio (WHR) but reduced IQCH-AS1 expression in adipose tissues. *,p≤0.05. ****, p≤0.0001. In (**H**) bars represent ± standard error.

## Discussion

Our current work interpreted population genetics of CAD using epigenetics of 45 disease-relevant cell types. We showed that on average 1.04% of CREs from the cells contained CAD-SNVs and approximately 50% of CAD-SNVs were located within CREs or TF binding sites of the 45 cell types (Fig. 1). By GATES test and SNEEP analysis, we linked these CREs and TF binding sites with CAD-SNVs to 1,580 candidate causal genes for the disease, including 1,208 protein-coding and 372 ncRNA genes. CAD-associated ncRNA gene showed better cell type specificity than the protein-coding counterpart (Fig. 2D). 792 of our candidate genes were also found by published studies or eQTL-based methods ^1–3,40^ and 798 were novel (Fig. 2). 542 out of 1,580 genes were regulated by the 38 TFs whose bindings were regulated by CAD-SNVs (Fig. 3). Both the novel protein-coding and ncRNA candidate genes showed similar PheWAS-eQTL colocalization patterns as the known candidates in terms of the associated traits and eQTL tissue types (Fig. 4), suggesting the reliability of our novel CAD candidate genes from epigenetics-GWAS integration. Our biological validation on one of the novel lncRNA genes, *IQCH-AS1*, indicated that the loss of the lncRNA could lead to detrimental adipose function, which was in line with its association with BMI and WHR, its downregulation in adipose tissue of CAD patients and the eQTL correlation in the same tissues (Fig. 5).

Our work identified more candidate causal genes (N = 1,580) for CAD than the published studies (N = 1,286) or eQTL-based methods (N = 1,169)^1–3,10,40^. Both the datasets and the statistical methodologies differ our work from others, which could lead to improved discovery. First, we included epigenetic data from 45 cardiovascular cell types, representing the most diverse cell groups investigated in CAD genetic studies. Second, our analyses also appear to be more sensitive than the GWAS-eQTL colocalization using GTEx and STARNET datasets, which identified 1,095 genes of nine CAD-relevant tissues in total. Given these eQTLs were mapped using tissue-level data, the eQTL signals with small effect sizes due to high cell specificity might be obscured in the bulk transcriptomic profiling. While future systematic single-cell eQTL datasets could make up for the drawback, our integration of cell type epigenetics has unveiled CAD candidate genes and CAD-TFs with high cellular specificity, such as *FLT1* in dendritic cells and *SLC22A3* or the AP-1 family TFs in adipocytes. Third, we used two complementary strategies, the GATES test and the SNEEP analysis, to prioritize CAD candidate genes. The GATES test summarizes the significance of all SNVs in the gene region and thus considers SNVs in coding and non-coding genic regions. The SNEEP analysis combines the prediction of TF-SNVs with the occurrence in cell type-specific regulatory elements and is thus focused on the non-coding part of the (epi)genome. That these tests draw their power from different statistical principles explains their complementarity. It is noteworthy that the way the prioritization is done using SNEEP is more stringent than in a previous application^41^. A controlled randomization approach suggested the use of the occurrence of at least *two* regulatory elements with TF-SNVs not in LD as a more stringent threshold to avoid false positives (Supp Fig. 1C). Fourth, genetically regulated non-coding RNA genes were systematically explored, which increased our discoveries. Among our CAD candidate genes, 23.5% were non-coding RNA genes, representing the highest ratio compared to reported candidate genes and genes found via GWAS-eQTL colocalization analysis.

For the first time, microRNAs were prioritized as candidate genes for CAD by genetic analysis. This was otherwise not possible by the population transcriptome-based method, given the existing genotyped transcriptome datasets, such as GTEx and STARNET datasets, excluded short RNA from the profiling. Interestingly, microRNA132 (MIR132), a therapeutic target in a Phase 2 clinical trial (NCT05350969) for heart failure^22^, was also exclusively underlined by our analysis (Supp Fig. 1E). Our analysis suggested MIR132 might also play a role in CAD. Indeed, the CRE SNVs for this ncRNA gene were not only from cardiac cells but also immune cells. CRE SNVs of MIR132 in immune cells could contribute to CAD. In addition, 361 ncRNA coding genes were prioritized, among them only 14 were conserved between humans and mice, which permits biological studies and therapeutic explorations both *in vitro* and *in vivo*. In our functional validation experiments, we demonstrated the novel ncRNA, *IQCH-AS1*, indeed was involved in CAD-relevant cellular functions.

Our diverse datasets and bioinformatic analyses might guide biological studies on gene functions. The GATES test and SNEEP analysis using the latest CAD GWAS summary statistics and epigenetic data from 45 cardiovascular cell types allowed pinpointing the candidate causal genes for the disease in a cell type-specific manner. The result will facilitate the selection of the relevant cell types to study candidate genes of interest. Our SNEEP analysis further indicated whether a TF could be involved in regulating the candidate gene expression in CAD. Finally, based on the PheWAS analysis, disease-relevant cellular phenotype(s) related to the candidate genes could be explored in the selected cell type(s). For instance, *IQCH-AS1* was regulated by the CAD-SNVs mediated epigenetic changes in stromal cells, mesenchymal stem cells, and adipocytes, all of which could affect adipose tissue function and cardiovascular health. Further, our PheWAS analysis indeed suggested *IQCH-AS1* to be associated with BMI and WHR (Fig. 4D&4E). Therefore, as proof of concept, we knocked down *IQCH-AS1* in preadipocytes and adipocytes and showed that the gene affected preadipocyte proliferation and lipid accumulation and therefore could play roles in obesity and CAD. Further molecular mechanisms shall be examined to explore the therapeutic potential of this RNA.

Limitations exist in our study. First, our analysis can not differentiate the role of the microRNA genes from the host genes. We therefore excluded microRNA genes in intragenic regions from our final candidate gene list. Second, the collection of epigenome data consists of different assays (DNase-seq, ATAC-seq, H3K27ac ChIP-seq) which can differ in the regulatory regions they uncover and in turn determine which SNVs can be found via epigenome analysis. However, a uniformly assayed collection for such a large collection of CAD-relevant cell types is not available. Third, the predictions for CRE-gene interactions allow one CRE to be linked to multiple genes. In these cases, we assume all target genes to be affected, although the effect might be limited to individual genes. We analyzed TFs whose binding sites are affected independently, despite TFs also functioning in a combinatorial manner^42,43^. Systematic incorporation of TF cooperativity would require knowledge of all possible TF interactions, which is not available. The prediction of TF-SNVs is based on TF binding motifs, which are imperfect for annotating binding sites, but enable us to analyze such a large collection of cell types and TFs.

Nevertheless, our study created an inventory of CAD candidate genes regulated by genetically affected epigenetics in a cell type-specific fashion. We prioritized novel candidate causal genes beyond existing methods, including protein-, ncRNA- and microRNA-coding genes. We demonstrated the significance of TFs in genetically regulated CAD risk. The results could lay the foundation for further biological investigation and the therapeutic exploration of CAD candidate genes.

## Methods

### Collection of enhancer-gene interactions

Epigenome data indicating CREs (DNase-seq, ATAC-seq, and H3K27ac ChIP-seq) were collected for 45 cell types related to CAD (Supp. Table 1&2). For each cell type the set of candidate CREs was defined as the peaks of the epigenome data. Most studies already provided peak annotations. For the scATAC-seq data of Hocker et al.^44^, the bigWig files of the cell types were converted to BedGraph with UCSC’s bigWig-ToBedGraph executable^45^, followed by MACS2’s bdgpeakcall function (v2.2.7.1)^46^ with a minimum length of 100 bp and signal cut-off of 0.004. The activity of the peaks was afterward assessed by taking the average signal of the bigWig-files. For the peak files of all cell types, replicates were merged and if the data was in genome version hg19 they were lifted to hg38 with a Python implementation of UCSC’s liftOver tool (v.1.1.13)^47^. Handling of bed-files was done with pybedtools (v0.8.1)^48,49^. To predict CRE-gene interactions for each cell type the generalized ABC-score from the STARE framework (v1.0.4)^15^ was used, using the average Hi-C data from Gschwind et al. as chromatin contacts^50^. The maximum distance between CRE and a gene’s TSS was set to 2.5 MB. CREs overlapping regions known to accumulate anomalous signals from sequencing experiments were excluded^51,52^. All interactions surpassing a gABC-score of 0.02 were considered relevant. The command was as follows:

STARE_ABCpp -b <peak_file> -n <activity_column> -a gencode.v38.annotation.gtf -w 5000000 -f <ENCFF134PUN_avgHiC_hg38/> -k 5000 -t 0.02 -x hg38-blacklist.v2.bed -o <output_path>.

### Identification of TF-SNVs and CAD-TFs

The GWAS summary statistic was downloaded from a recently published study for coronary artery disease^1^. Similar to Aragam et al., a 1% FDR cutoff was applied to extract those SNVs significantly associated with CAD (*p* ≤ 2.52E-5). SNVs in linkage disequilibrium (LD) were determined based on the pre-computed dataset from SNiPA^53^, where the European cohort and an LD threshold ≥ 0.8 were used. The lead and proxy SNVs were lifted to hg38 using the Python package liftover^47^. The resulting 72, 432 SNVs are used in the following analyses (Supp. Table 3).

The SNEEP software (version v1.0)^14^ was applied for the identification of TF-SNVs (Supp. Table 4), disease genes, and disease-associated TFs. Based on the beta coefficient given in the summary statistic (denoted as beta) the alleles were ordered in such a way that SNEEP compares the CAD-risk allele against the non-risk allele.

TF-SNVs were predicted for each of the 45 cell types individually. From the 72,432 lead and proxy SNVs, those not overlapping with the cell type-specific enhancer-gene interactions were excluded. Further, these interactions were used to associate the TF-SNVs to cell type-specific target genes. As TF motifs, 817 non-redundant human motifs from JASPAR (version 2022)^54^, HOCOMOCO^55^, and Kellis ENCODE motif database^56^ were used (publicly available at: https://github.com/SchulzLab/SNEEP/).

Further, 500 background analyses were performed based on randomly sampled SNVs. Whenever possible the random SNVs were derived with SNPsnap^57,58^ to ensure random SNVs were most similar to the original ones in terms of minor allele frequency (MAF), distance to the nearest gene, gene density, and SNVs in LD. However, for less than 5% of all lead and proxy SNVs, SNPsnap failed since the LD structure of the SNV could not be derived. In such a case the corresponding random SNV with a similar MAF was sampled from the dbSNP database^59^. The following SNEEP command was performed per cell type:

differentialBindingAffinity_multipleSNPs -p 0.5 -c 0.001 -f <celltypeSpecificEnhancerGeneinteractions_merged> -r <celltypeSpecificEnhancerGeneinteractions> -g <geneId_geneName.txt> -j 500 -l <seed> <combined_motif_set> <snvFile> <hg38.fa> <estimatedScalesPerMotif_1.9.txt>.

To explore genes significantly affected by the analyzed SNVs, the randomly sampled SNVs were linked to target genes using the cell type-specific enhancer-gene interactions. Next, it was counted per gene how many non-LD TF-SNVs in different CREs were observed in the random data. Based on the resulting cell type-specific background distribution over all genes, a FDR corrected p-value was derived. Overall cell types observing at least two non-LD TF-SNVs were significant (FDR ≤ 0.01), suggesting that the associated target genes are highly affected by the CAD-SNVs (Supp. Fig. 1C).

Based on the background analysis, 38 cell type-specific TFs were derived that are more often affected by the analyzed SNVs than expected on a random background control (Supp. Table 7). Therefore, for each TF an odds ratio was computed given the numbers of how often a TF’s binding site was significantly affected by the analyzed SNVs in comparison to the random SNVs. The analysis was done as described by Baumgarten et al.^14^ (see Star Methods Section, eQTL analysis of this paper). CAD-TFs are defined as those TFs with more than five TF-SNVs, an odds ratio ≥ 5 in at least one of the analyzed cell types, and an open promoter region in the same cell type.

### Functional consequences of CAD-SNVs with VEP

For the functional annotation of variants the Variant Effect Predictor (VEP) from Ensembl (v111.0)^11^ was used with the ‘pick’ option to get one consequence per variant. The distance to consider up- or downstream variants was set to zero. Categorical colors were taken from the colorcet Python package, based on Glasbey et al.^60^.

### Gene-based test

In addition to SNEEP the gene-based association test GATES^13^ was applied to identify CAD-associated genes. For each gene, all SNVs from the CAD GWAS summary statistic^1^ (lifted to hg38) that overlap the genes’ bodies were considered. The gene body was defined as annotated in the GENCODE’s^61^ v38 gtf-annotation including exons and introns. To determine the LD structure for all pairs of SNVs associated with a gene, PLINKs^62^ functionality (http://pngu.mgh.harvard.edu/purcell/plink/, version 1.07) to look up pair-wise LD correlation (R2) was used:

plink-1.07-x86_64/plink --noweb --bfile alkesgroup.broadinstitute.org/LDSCORE/GRCh38/plink_files/<chromosome> --snps <list_rsids> --r2 --matrix --ld-snp-list <gene> --out <gene>.txt.

The LD score data from PLINK was downloaded from https://alkesgroup.broadinstitute.org/LDSCORE/GRCh38/ in March 2022. The SNVs for which the pair-wise LD score could not be derived were excluded. Next, GATES was computed for each gene with SNVs in the gene body. The resulting p-values were FDR-corrected with the Benjamini-Hochberg procedure^63^. The implementation for GATES was taken from the R package COMBAT^64^ and slightly adapted.

### Defining candidate CAD genes and adding further evidence

We combined the results of the TF-SNV analysis and the gene-based association test to define a joint set of candidate CAD genes. All genes from the SNEEP analysis that had at least two non-LD TF-SNVs in different CREs in at least one cell type in which the gene was also active were considered candidate CAD genes. A gene was considered active in a cell type if a peak called on the epigenetic signal overlapped any of the gene’s promoters (± 200 bp, Supp. Table 5). For the GATES test, a gene had to have at least 10 SNVs in its gene body, be open in any cell type, and have an FDR ≤ 2.5%. Specific genes were excluded from all gene sets and not considered as potential candidate CAD genes, all based on GENCODE’s^61^ v38 gene annotation: pseudogenes, genes that are not yet experimentally confirmed (labelled as TEC), miRNAs that are located completely in the gene body of other genes on the same strand and genes which share their 5’ TSS with other genes. Overall, 17,732 genes from the annotation were excluded (Supp. Table 5).

Additional evidence and data were gathered for the candidate CAD genes (Supp. Table 5). It was checked whether the candidate genes were also identified in other types of analyses. A manually curated list of known CAD loci genes was generated, comprising 1,286 genes (after removing the excluded genes) and which is based on previously published work combining the following criteria: 1) the biological plausibility, 2) rare coding variant(s) associated with CAD, 3) likely pathogenic variant(s) relevant to CAD in ClinVar^5^, 4) evidence from effective cardiovascular (CV) drug(s), 5) significant causality by Mendelian Randomization studies, 6) a protein-altering variant in high LD with the sentinel CAD variant, 7) expression quantitative trait loci (eQTLs) in a CAD-relevant tissue, and 8) a CV-relevant phenotype in knockout mouse models^1,2,6^. Gene names were mapped to Ensembl IDs with the help of my MyGene.info ^65–67^.

Further, it was added whether a gene was also found via colocalization analysis with eQTL data from GTEx^17^ and the STARNET studies^18^ or among candidate CAD genes from a recent publication that focused on endothelial cells^10^. In addition, it was tested whether the candidate CAD genes are co-expressed with known CAD loci genes. To do so, a co-expression analysis was performed using gene expression profiles of 9,662 GTEx RNA-seq samples^17^. For each candidate CAD gene, the top 100 co-expressed protein-coding genes were derived using the Spearman correlation coefficient as a similarity metric. Applying Fisher’s exact test, 74 candidate CAD genes were significantly co-expressed with known CAD loci genes (FDR ≤ 0.05). The expression breadth was determined based on RNA-seq data from the IHEC EpiATLAS (1,555 samples, https://ihec-epigenomes.org/epiatlas/data/) and calculated as the fraction of cell types/tissues (N = 58, metadata column ‘harmonized sample ontology intermediate’, metadata version 1.1) in which a gene had an expression of ≥ 0.5 transcripts per million. An expression breadth of 1 means that a gene is expressed in all cell types/tissues. If a gene is conserved in mouse was based on the annotated orthologues from BioMart^68^ (confidence > 0). To also account for lncRNAs, a Reciprocal Best BLAST Hit^69^ was performed (sequence version GRCm39 for mouse and GRCh38 for human). The human lncRNA sequences were aligned to the mouse genome (forward) and the process was repeated the other way around (reverse). If a human gene had a reciprocal best match without duplicate alignments, it was considered to be conserved in mice. Overlap of gene sets were visualized with UpSet plots^70^. All GO term enrichment tests were done with g:Profiler’s Python package (v.1.0.0)^71^. For the enrichment test of the candidate CAD genes (Fig. 2B), all genes active in any cell type and not part of the set of excluded genes were used as background. As background for the TF target genes, all genes active in any cell type were used.

### Differential expression of CAD-TFs in CAD patient data

The differential expression data for the atherosclerotic aortic root, liver, subcutaneous fat, skeletal muscle, and visceral fat tissues between 600 individuals with CAD and 250 CAD-free controls were estimated using the DESeq2^72^ R package and sourced from the supplementary materials of the STARNET consortium study^18,73^ (Supp. Table 8). Genes with an adjusted p-value of < 0.01 and an absolute log2FC > 0.3 are defined as differential. 34 out of 38 CAD-TFs were available in the STARNET data set (Fig. 3B&3C). The expression of the target genes (log2FC) of TFs TFAP2E, ZIC4, ZBTB33, MBD2, ZNF610, TCFL5, and ZFP14 was extracted for the tissues in which the TF was differentially expressed. Using a one-sided Wilcoxon-rank sum test (R function Wilcox. test, alternative = greater), it was derived whether the target genes expression differs between the tissues. Significant differences (p-value ≤ 0.05) are labeled with an ∗ in Figure 3E.

### eQTL support for CAD-genes

The eQTL data for 49 tissues and cell types was downloaded from the GTEx^17^ portal (version 8). Only the eQTLs linked to known CAD GWS genes or candidate CAD genes were kept. Additionally, it was filtered whether the eQTL was found in a CAD-relevant tissue included in the GTEx data (aorta, coronary artery, mammary artery, tibial artery, visceral and subcutaneous adipose tissues, whole blood, heart atrial appendage and left ventricle, kidney cortex, liver, skeletal muscle, spleen, and tibial nerve). For each CAD-TF, the TF-SNVs from the cell types the CAD-TFs were identified in were gathered. The fraction of eQTLs supporting the TF-SNVs linked to target genes was computed (Fig. 3D right panel, Supp. Table 9).

### GWAS sources and eQTL datasets for colocalization analysis

The 29 GWAS datasets of CAD-relevant traits or diseases were full-genome summary statistics from publicly available studies (Supp. Table 12). eQTLs of artery (aorta, coronary, mammary, tibial), adipose tissues (visceral, subcutaneous), blood, heart (atrial appendage, left ventricle), kidney cortex, liver, skeletal muscle, spleen, and tibial nerve, were obtained from STARNET^18^ and GTEx^17^ v8 datasets. The STARNET dataset was obtained via collaboration and the GTEx dataset was accessed via the dbGaP platform.

### GWAS-eQTL colocalization analysis with COLOC

COLOC represents a Bayesian approach that estimates the posterior probabilities between a given GWAS signal and a given QTL^16,74–76^:

H0: Neither has a significant association in the region

H1: Only the GWAS trait has a significant association in the region

H2: Only the QTL trait has a significant association in the region

H3: Both GWAS and QTL traits have a significant association in the region, but the variants are different

H4: Both GWAS and QTL traits have a significant association and share the variants in the region

For each CAD candidate gene, we obtained eQTL SNVs from GTEx and STARNET studies with p-value < 0.01. These SNVs were used to test the colocalization with the GWAS associations of the 29 datasets. GWAS SNVs with p-value < 5e-8 were selected for the analysis. To estimate the posterior probability of GWAS-eQTL colocalization we used the coloc R package and ran the coloc.abf function. GWAS and eQTL datasets with the effect sizes, allele frequencies, and sample sizes were used as inputs. Significant colocalizations were defined by PPH4 ≥ 0.60, indicating shared association signal and variants between GWAS and eQTL (Supp. Table 11). Top genes were selected based on their GWAS association strength, with priority given to the lowest GWAS p-values and the highest number of colocalized traits.

### Mapping cis regulated eQTL and gene expression in adipose tissue

RNA sequencing and blood genotyping for STARNET subcutaneous fat (SF) and visceral fat (VAF) tissues were performed as described^18,73^. Bulk RNAseq and genotyping data were processed for quality control, genotyping data imputation, expression normalization, and cis-regulated expression quantitative trait loci (eQTLs) inferral^18,73^. Results of cis regulated eQTL for SNV rs3784699 and *IQCH-AS1* gene expression in SF and VAF were plotted using ggplot2 in R.

### Cell culture and adipogenic differentiation

Human Simpson-Golabi-Behmel Syndrome (SGBS) preadiocytes, a non-immortalized preadipocyte cell line, were provided by Dr. Daniel Tews and Prof. Martin Wabitsch, and cultured in DMEM/F-12 (#11330057, ThermoFisher, Waltham, USA) supplemented with 10% FBS (#S0615, Sigma Aldrich, St. Louis, Missouri, USA) and 100 U/l penicillin/streptomycin (#15140122, ThermoFisher, Waltham, USA) at 37 °C in 5% CO_2_. Cell differentiation was performed as described previously^77^.

### CRISPR/Cas9 plasmids and virus infection

SgRNA sequences for lentiviral plasmid construction were designed using the online tool available at^78^. Two complementary oligonucleotides were synthesized: 5’-CACCG-[sgRNA sequence-3]’ and 5’-CAAA-[reverse complement of sgRNA]-C-3’. As described previously^79^, two sgRNAs targeting the shared exon of all transcripts were delivered via lentivirus into SGBS-preadipocytes. Exon 13 of IQCH-AS1 was specifically targeted using a dual CRISPR strategy, resulting in a 48 bp frameshift deletion. The sgRNAs were carried by the LentiCRISPR v2 (#52961, Addgene, Watertown, USA), and cells were selected with 2 μg/mL puromycin for 5 days to remove non-infected cells. The positively targeted cells were then expanded in culture and subsequently passed for further analysis.

### Amplification polymerase chain reaction (PCR) and Real-time PCR(RT-qPCR)

Total RNAs were isolated from cells using TRIzol reagent, and an amount of 3 µg of RNA was used in 10 μl reaction volume to digest DNA using the Maxima H Minus cDNA Synthesis Master Mix (#15606029, ThermoFisher, Waltham, USA) following the manufacturer’s instructions. PCR amplification was performed with Q5 High-Fidelity 2X Master Mix (#M0492L, New England Biolabs, Massachusetts, USA) using 750 ng of cDNA template on a thermal cycler. The PCR products were electrophoresed using 1% agarose in 1×TBE buffer at 120 V for 45 min, with *GAPDH* as an internal control (Supp. Table 13). For RT-qPCR, SYBR Green probes (#95074-012, VWR International, Radnor, Pennsylvania, USA) were used to detect the target genes and 60 ng cDNA was used as the template. *GAPDH* was used as an internal control. Reactions were performed on a ViiA 7 Real-Time PCR System (ThermoFisher, Waltham, USA). Expression levels were reported as 2-DCt values (Supp. Table 13).

### Lipid extraction and triglycerides measurement

As described previously^79^, the SGBS adipocytes were first washed twice with PBS, and then detached by scraping. Lipids were extracted using a chloroform mixture (2:1, v/v), followed by overnight drying in a fume hood. The dried lipid residue was resuspended in 100 μL of 1% Triton X-100 in absolute ethanol and incubated for 1 hour with constant rotation. After incubation, the suspension was dried in a SpeedVac for 30 minutes, the residue was then resuspended in 100 μL of PBS with 1% Triton X-100. To measure lipid content, a 3 μL aliquot of the suspension was taken, and lipid triglycerides were measured using a triglyceride determination kit (#TR0100, Sigma Aldrich, St. Louis, USA).

### Immunofluorescence staining

Cells were washed with PBS and fixed in ice-cold methanol at −20 °C for 15 min. After PBS washing, cells were blocked for 1h at room temperature with PBS containing 5% m/v BSA. Cells were incubated overnight at 4 °C with rabbit anti-PPARgamma primary antibody (1:200 dilutions) (#2443S, Cell Signalling Technology, Danvers, USA). After PBS washing, Donkey anti-rabbit lgG Highly Cross-Adsorbed Secondary Antibody (#a32790, ThermoFisher, Waltham, USA) and HCS LipidTOXred neutral lipid stain (#H34476, ThermoFisher, Waltham, USA) were applied. Cells were then mounted with a DAPI mounting buffer (#62248, ThermoFisher, Waltham, USA). Fluorescence signals were visualized by the THUNDER imaging system (Leica, Wetzlar, Germany).

### Cell proliferation assay

The BD Pharmingen™ APC BrDU Kit (#552598, BD Biosciences, Franklin Lakes, USA) was used following the manufacturer’s protocol. Cells were treated with 10 μmol/L BrdU for 24 hours, then were collected, fixed, and permeabilized using the kit. BrdU incorporation was assessed with a BD LSRFortessa™ Cell Analyzer (BD Biosciences, Franklin Lakes, USA). The recorded events were gated using forward- (FSC-A) and side-scatter (SSC-A) adopted to the cell type. Then, single cells were gated, using FSC-A versus forward scatter height (FSC-H). Finally, within the single cell population, BrdU^+^ population was identified. Unstained control samples were used as negative control of the fluorescence and to set the gate BrdU^+^ cells. Gating was applied to exclude doublets and flow cytometry data were analyzed using FlowJo (version 10.8.2, RRID: SCR 008520).

### Proteome Profiler Human XL Cytokine Array

Cytokines released from control and IQCH-AS-KO adipocytes were analyzed using the Proteome Profiler Human XL Cytokine Array (R&D Systems, Minneapolis, MN, USA). Specifically, cells for the assay were cultured for 24 h in a serum-free medium. Mediums were first collected and enriched five times by vacuum centrifuge, then incubated with a biotinylated detection antibody cocktail on a nitrocellulose membrane. After washing, the membrane was incubated with streptavidin–horseradish peroxidase. Cytokine signals were detected with the ImageQuant 800 imaging system (Amersham Biosciences, Amersham, UK). Cytokine signal density was quantified using dot plot analysis in the ImageQuant TL software and then normalized by subtracting the background intensity. The normalized signal density was compared between control and *IQCH-AS1*-KO adipocytes.

### Statistical analysis

Normality of datasets, excluding cytokine arrays, was assessed using the Kolmogorov-Smirnov or Shapiro-Wilk test. Data conforming to a normal distribution were analyzed using two-tailed unpaired t-test. Results are presented as mean ± SEM. For the analysis of cytokine array data, multiple unpaired t-tests were employed. The false discovery rate was controlled at 5% using the Benjamini-Hochberg correction for multiple comparisons^63^. Statistical calculations were conducted using GraphPad Prism 10 (GraphPad Software, La Jolla, CA; RRID: SCR_002798).

## Acknowledgments

This work was partly supported through the computational and data resources and staff expertise provided by Scientific Computing and Data at the Icahn School of Medicine at Mount Sinai.

## Author contributions

Study design and supervision by M.H.S and Z.C. Collection of epigenome data, prediction of enhancer-gene interactions and related analyses by D.H. Running SNEEP and analyses on TF-level were done by N.B. GATES test by N.B. and N.K. Co-expression analysis by N.B. and F.B.A. Sequence similarity analysis by R.K.M. Co-localization analysis with eQTL data by L.L. and A.D. Collection of known CAD disease genes by Z.C. Integration of data and analyses on gene-level by D.H. PheWAS analysis and processing of STARNET expression data by A.D. Collaboration for STARNET data with J.L.M.B. and L.M. Biological experiments by X.S. and S.L., for which SGBS preadiocytes were provided by D.T. and M.W. Manuscript writing by D.H., X.S, N.B., A.D., M.H.S., Z.C. Feedback on manuscript all authors.

## Data availability

All data generated or analysed during this study are included in this published article (and its supplementary information files), except for the raw STARNET which was made accessible by members of the STARNET project.

## Code availability

SNEEP for predicting TF-SNVs is available under https://github.com/SchulzLab/SNEEP/. STARE for regulatory interactions is provided under https://github.com/SchulzLab/STARE. The software for the colocalization analysis was taken from https://cran.r-project.org/web/packages/coloc/index.html. Custom scripts used in this manuscript can be found on GitHub https://github.com/SchulzLab/CADLinc.

## Funding

The work was funded by the Sonderforschungsbereich SFB TRR 267 (DFG, 403584255, project B05 ZC & HS and project Z03 MHS), the German Research Foundation (DFG 510049865) to ZC, Corona-Stiftung Junior Research Group Grant to ZC and the Federal Ministry of Education and Research (Bundesministerium für Bildung und Forschung, BMBF) as part of the German Center for Child and Adolescent Health (DZKJ) under the funding code 01GL2407A to MW and DT. We acknowledge the support of the German Centre for Cardiovascular Research (DZHK) with a Postdoc Start-up Grant (ID: 81X3600510) to ZC, the DZHK (IDs: 81Z0200101 and 81X2200151) to MHS, the Cardio-Pulmonary Institute (CPI) [EXC 2026] ID: 390649896 to MHS, the DFG SFB1531 (S03, project ID 456687919) to MHS, and the Clinical and Translational Science Awards (CTSA) grant UL1TR004419 from the National Center for Advancing Translational Sciences to LM.

## Notes

https://github.com/SchulzLab/CADLinc

### Competing Interest Statement

The authors have declared no competing interest.

