## Supplementary figures for "Cell type-specific epigenetic regulatory circuitry of coronary artery disease loci"

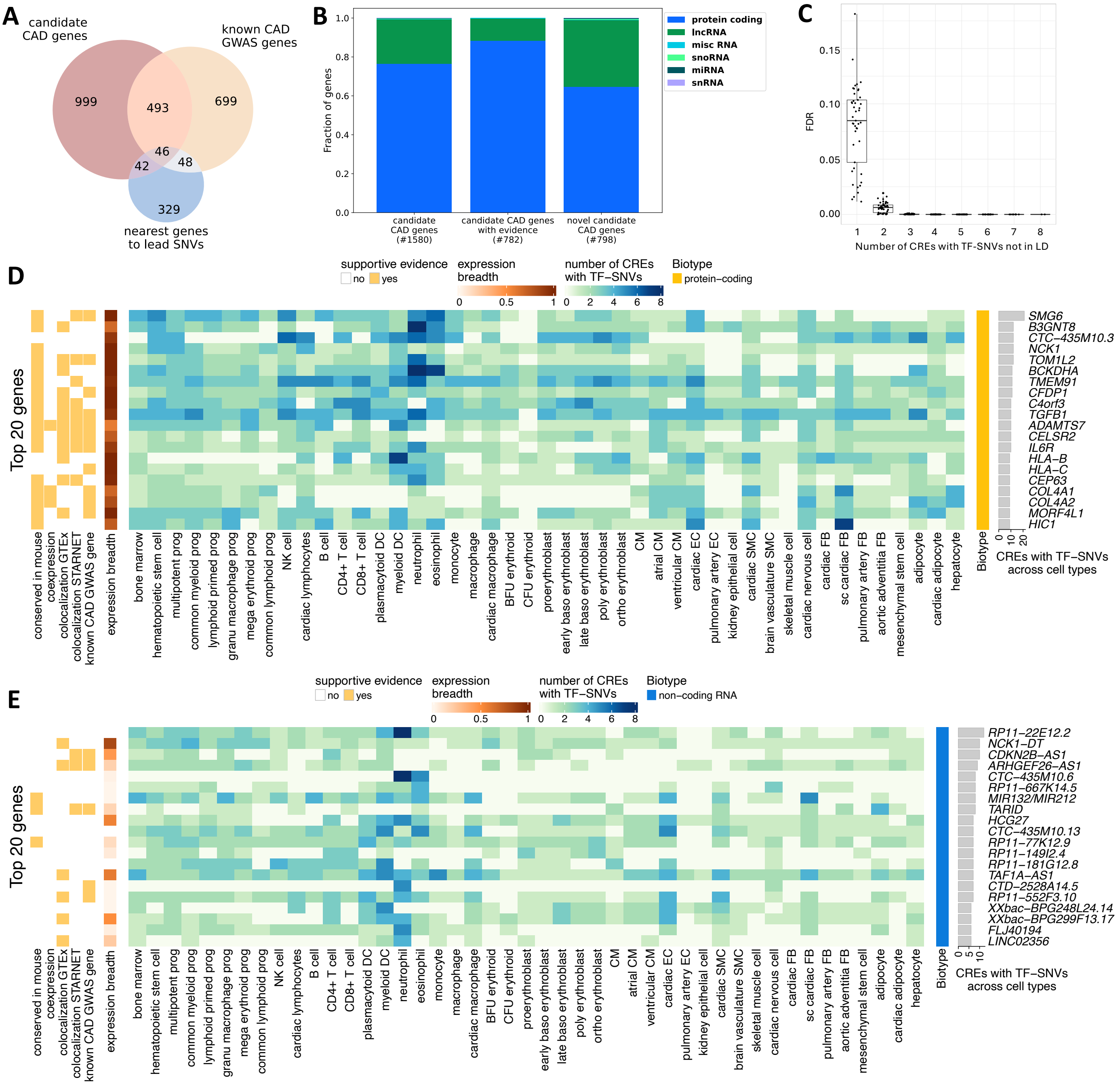


**Supp. Figure 1**: **Extended information on candidate CAD genes. (A)** Venn diagram of our candidate CAD genes, known CAD GWAS genes and gene that are closest to 688 lead CAD SNVs. Only genes with epigenetic promoter signal in any cell type and not among the excluded genes were considered. **(B)** Biotypes of all candidate genes, the candidates with previous evidence (based on literature and colocalization analysis) and the candidates that are novel. The number in parentheses indicates the size of the gene set. **(C)** Boxplot that shows how likely it is to observe a certain number of CREs with TF-SNVs not in LD to each other on randomly sampled SNVs. The analysis was performed for each cell type separately (dots). SNVs are sampled using SNPsnap[57,58] to assure similar properties as for the CAD GWAS SNVs within the cell type-specific CREs (see Methods 'Identification of TF-SNVs and CAD-TFs'). **(D-E)** Heatmap of the top 20 protein-coding **(D)** and non-coding RNA genes **(E)** ranked by the number of merged CREs with TF-SNVs, analogue to Fig. 2G.

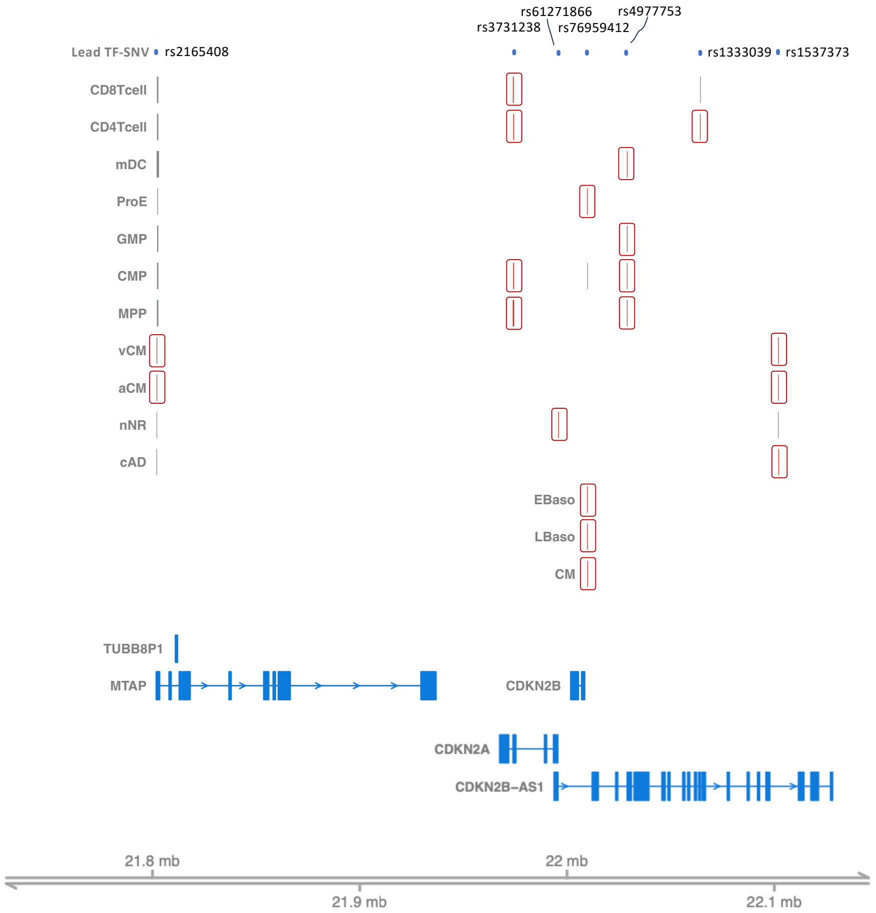


**Supp, Figure 2**: **Detailed overview of the CDKN2B-AS1 loci**. Lead TF-SNVs (upper row, blue dots) and their overlapping CREs (black and red bars) from different cell types are visualised. Red marked CREs are linked to CDKN2B-AS1.

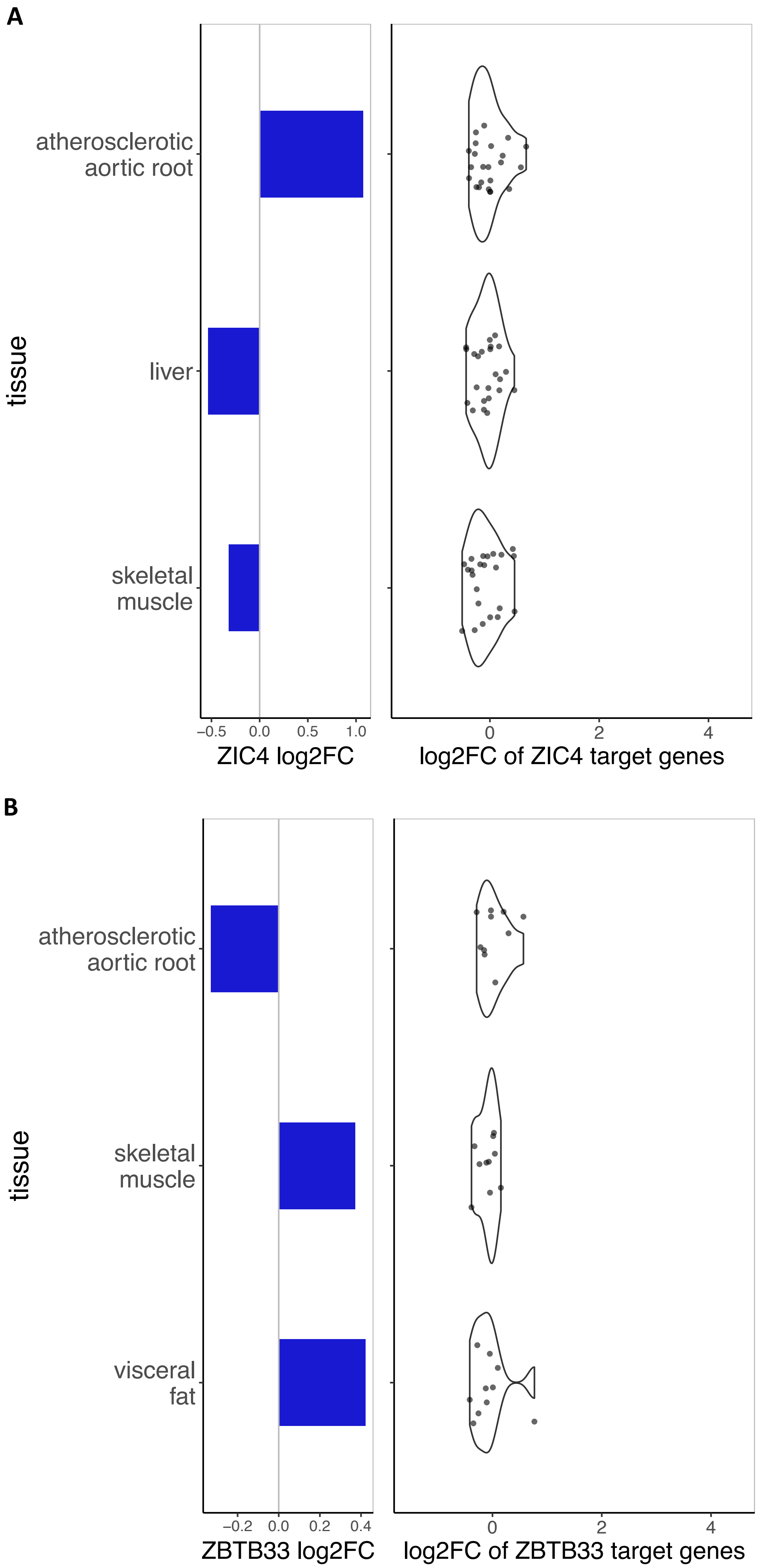
A
tissue
atherosclerotic aortic root
liver
skeletal muscle
−0.5 0.0 0.5 1.0
ZIC4 log2FC
0 2 4
log2FC of ZIC4 target genes
B
tissue
atherosclerotic aortic root
skeletal muscle
visceral fat
−0.2 0.0 0.2 0.4
ZBTB33 log2FC
0 2 4
log2FC of ZBTB33 target genes

C

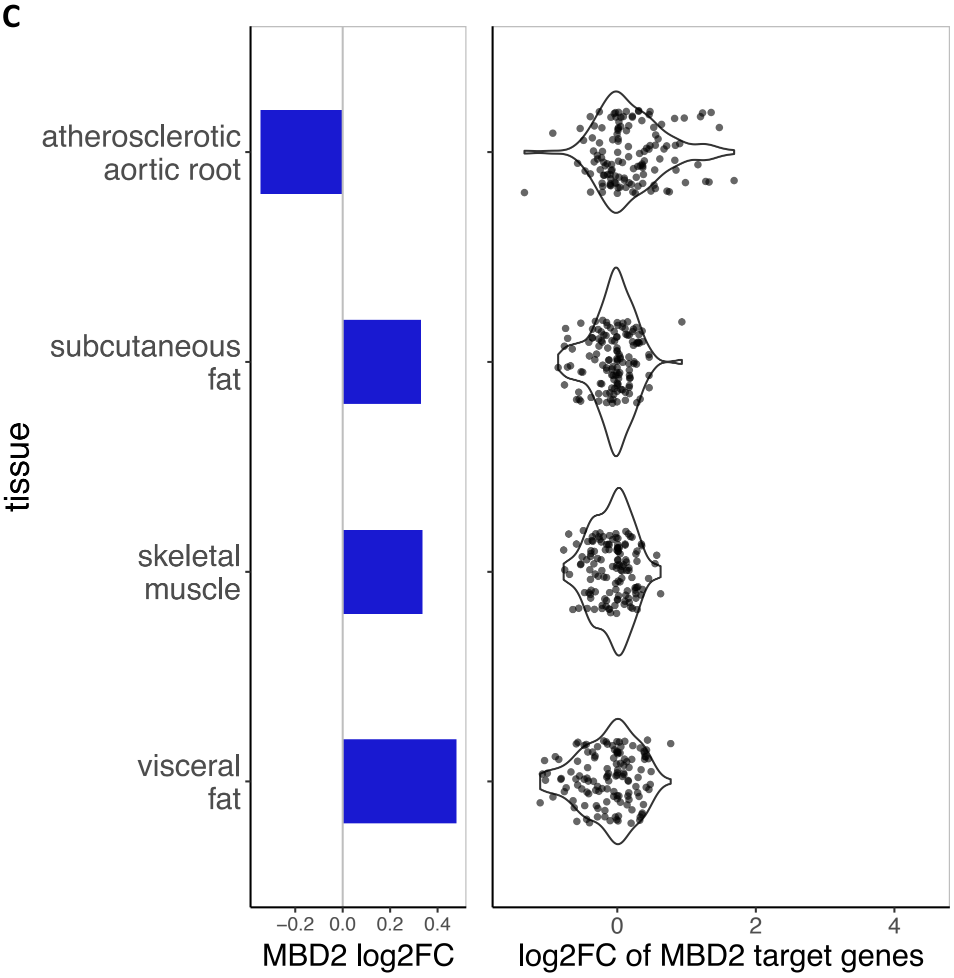
atherosclerotic aortic root
subcutaneous fat
skeletal muscle
visceral fat
tissue
−0.2 0.0 0.2 0.4
MBD2 log2FC
0 2 4
log2FC of MBD2 target genes

D

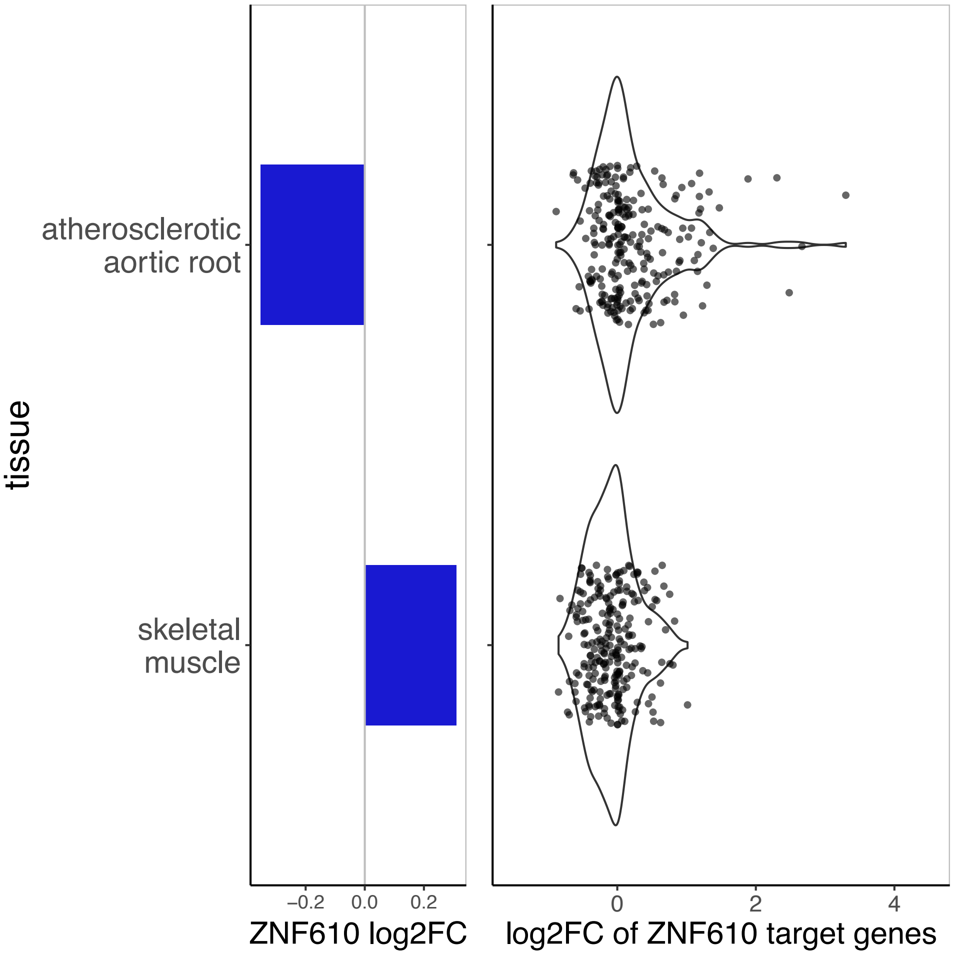
atherosclerotic aortic root
skeletal muscle
tissue
−0.2 0.0 0.2
ZNF610 log2FC
0 2 4
log2FC of ZNF610 target genes

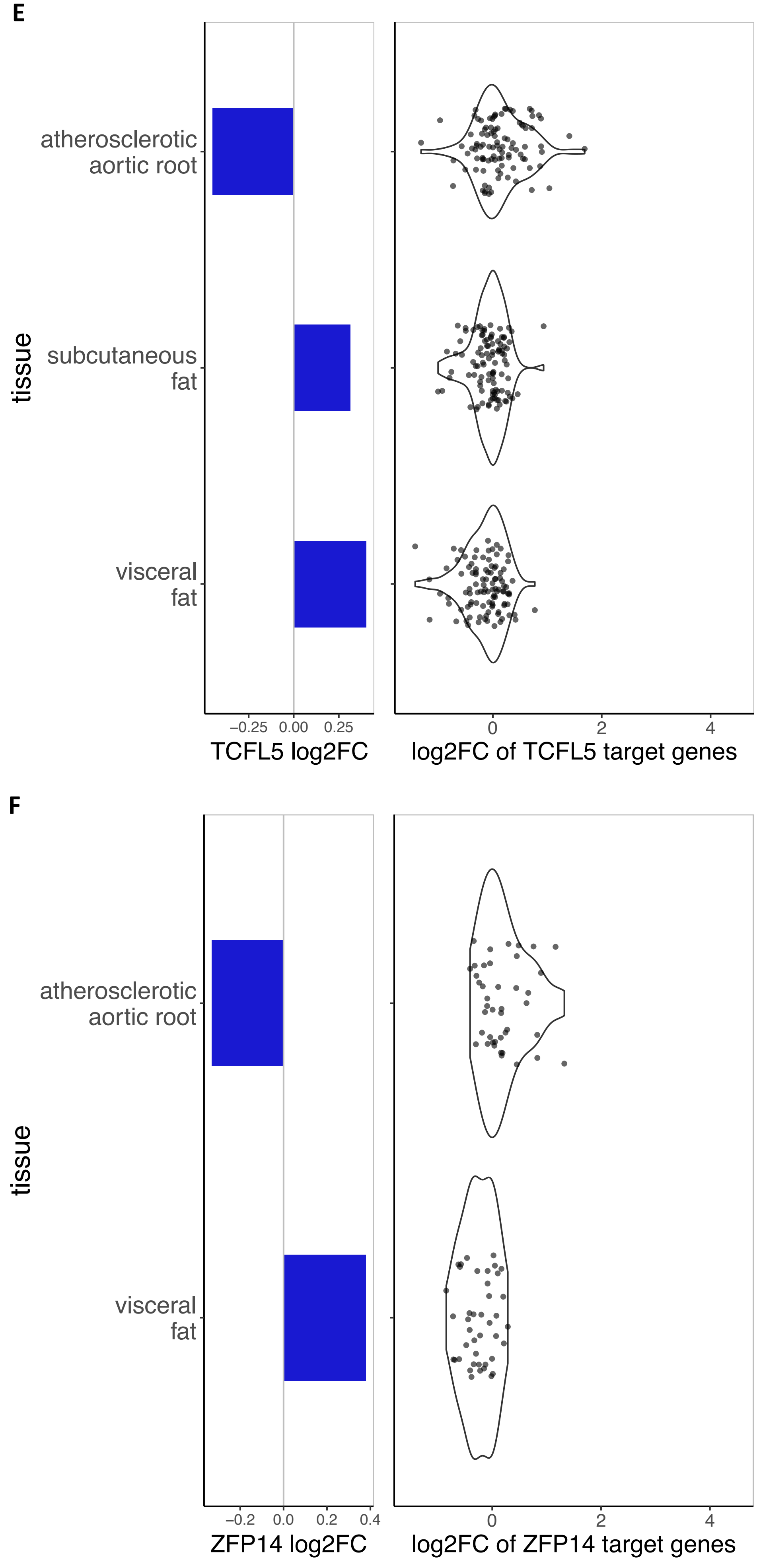


**Supp. Figure 3**: **Further examples similar to Figure 3E.** Analysis of TF gene deregulation (log2 fold change, barplot) and expression deregulation of associated TF target genes in different STARNET[18] tissues (log2 fold change, violin scatter) for **(A)** ZIC4, **(B)** ZBTB33 **(C)** MBD2 **(D)** ZNF610, **(E)** TCFL5 and **(F)** ZFP14. Only tissues in which the TF is expressed are included.

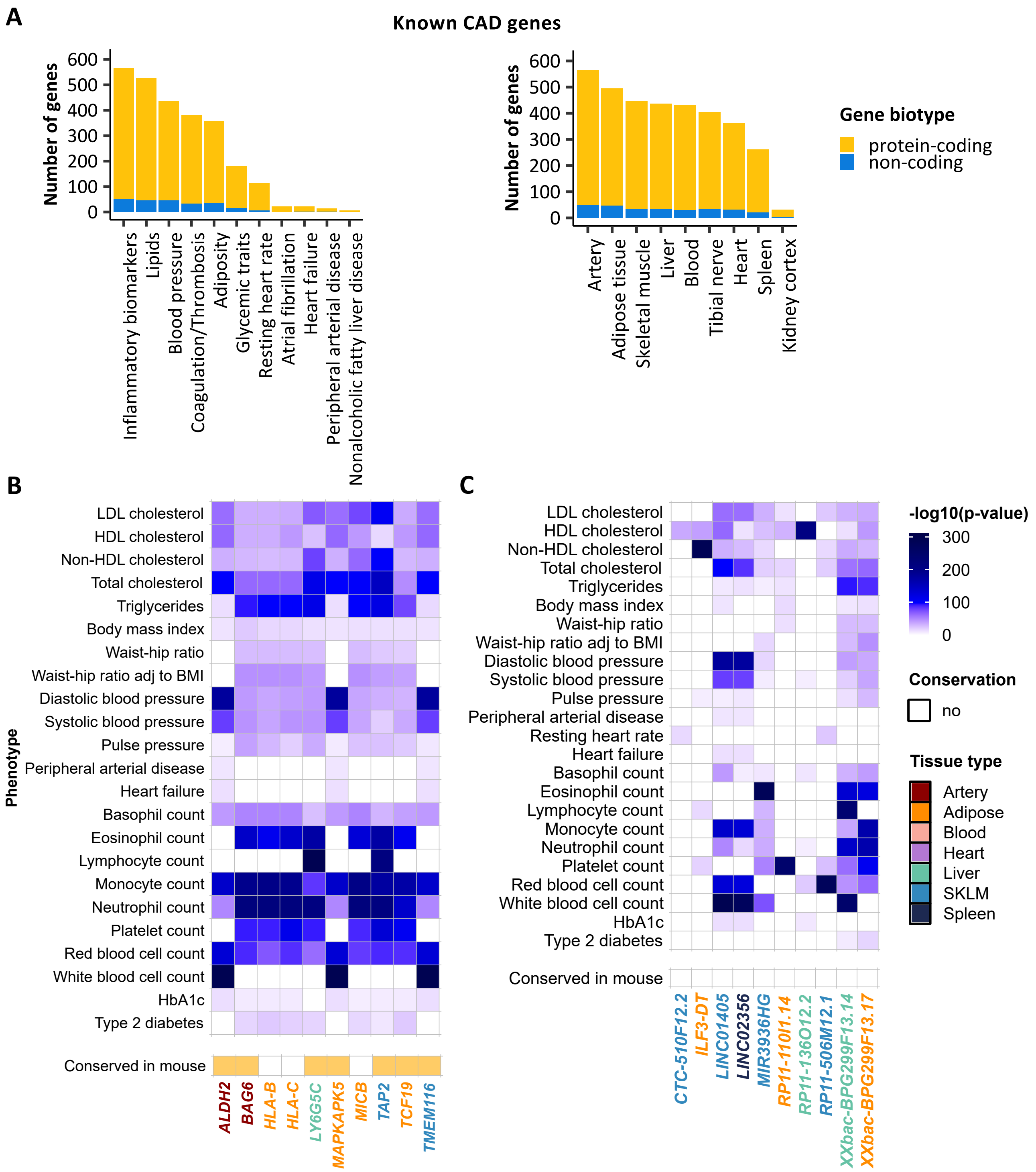


**Supp. Figure 4: PheWAS-eQTL colocalization analysis on known CAD genes among our candidate genes. (A)** Number of protein-coding and non-coding known CAD genes with significant colocalized GWAS-eQTL signals across phenotypes and tissue types. **(B-C)** Top 10 known CAD protein-coding **(B)** and non-coding **(C)** genes with the strongest eQTL effect on CAD-related phenotypes.

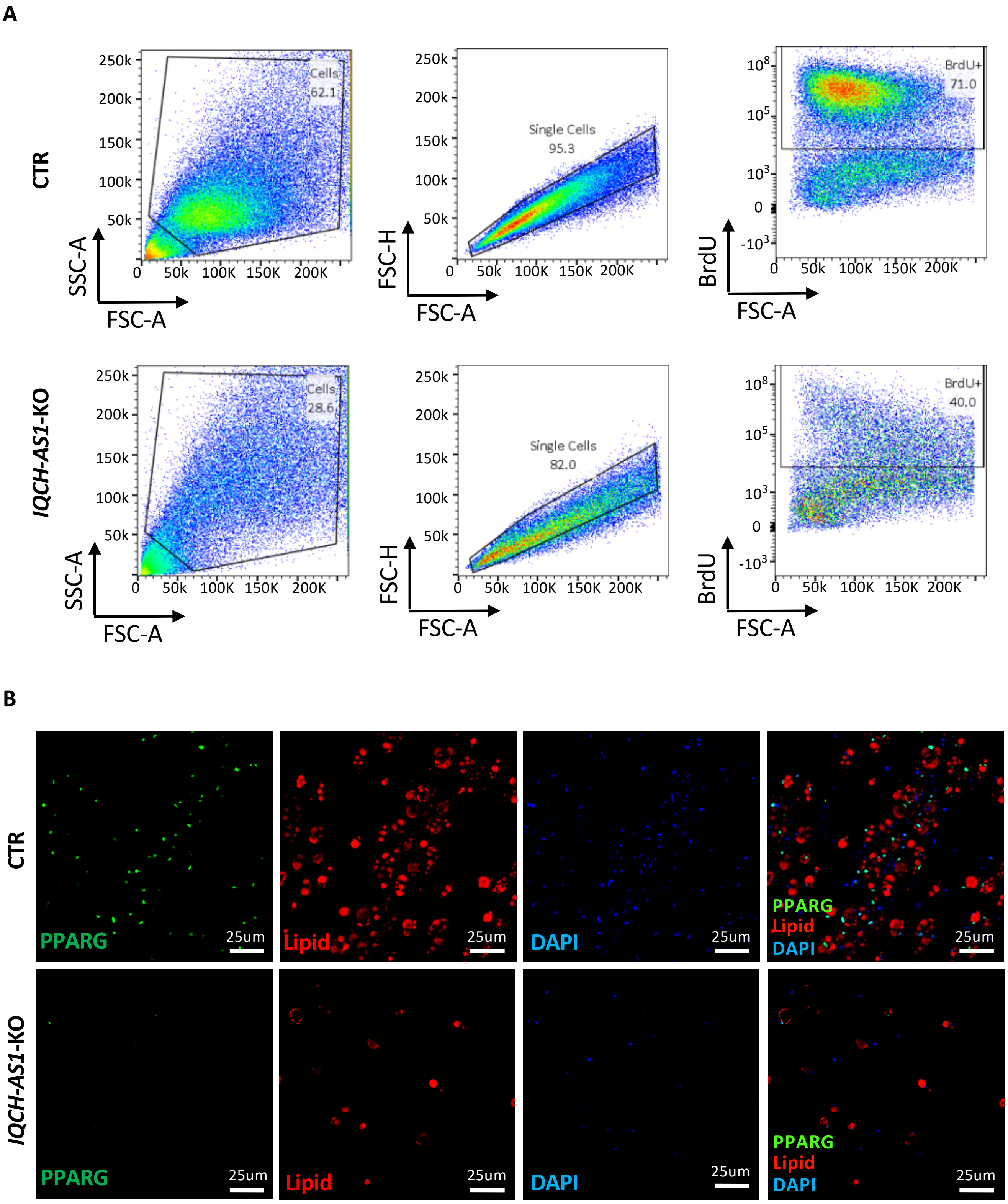


**Supp. Figure 5: Gating strategies and immunostaining.** (**A**) The gating strategies in control and knockout groups (full images of Fig. 5C). Immunostaining of control (CTR) and *IQCH-AS1*-KO adipocytes (**B**) (full images of Fig. 5D). PPARG was stained by the protein-specific antibody (green). Lipid droplets were labeled by HCS LipidTOX$^{TM}$ Red neutral lipid stain (red), and DNA by DAPI (blue).
