## Supplementary material for "Cell type-specific epigenetic regulatory circuitry of coronary artery disease loci": Table of supplementary files

Table with files in the Supplementary Material folder

| Name | Description |
|---|---|
| Supp.Table1-EnhancerMetadata.txt | Metadata for epigenome data for all 45 cell types |
| Supp.Table2-EnhancerInteractions_hg38.zip | Folder with REMs per cell type and their target genes. One bed-file per cell type with all candidate CREs and their comma-separated target genes in the 4th column. |
| Supp.Table3-GWAS_CAD_SNVs_hg38.txt | List of all CAD-SNVs (hg38) |
| Supp.Table4-SNEEPTable_hg38.txt | SNEEP table, contains which SNVs, TFs, CREs and genes (hg38) |
| Supp.Table5-CandidateCADGenes.xlsx | Gene table with all the integrated information and additional gene lists: known CAD GWAS genes, genes found via colocalization analysis, active genes, excluded genes |
| Supp.Table6-CandidateCADGenes_gProfiler.tsv.gz | Complete GO result table of the candidate CAD genes |
| Supp.Table7-TFTable_Fig3A.xlsx | TF table with enrichment across cell types and input to Fig. 3A |
| Supp.Table8-STARNET_DifferentialExpression.xlsx | CAD STARNET differential expression |
| Supp.Table9-TFSNV_eQTL_Overlap_Fig3D.txt | Overlap of TF-SNVs with eQTLs |
| Supp.Table10-CADTFs_TargetGenes_gProfiler.xlsx | Complete GO result table of CAD-TF target genes |
| Supp.Table11-PheWAS_eQTL_Colocalization.csv | Table with PheWAS and colocalization results |
| Supp.Table12-GWAS_Phenotypes.xlsx | List of GWAS sources with CAD-relevant phenotypes |
| Supp.Table13-Primers.xlsx | List of primers for PCR and RT-qPCR |
| Supp.Table14-IQCH_AS1_CREs_AD_MSC_stromal_hg19.txt | CREs of IQCH-AS1 in stroma, AD and MSC (hg19) |
| Supp.Table15-CDKN2B_AS1_CREs_Supp2_hg38.txt | CREs of CDKN2B-AS1 for Supp. Fig. 2 (hg38) |
